# Claudin-11 regulates immunological barrier formation and spermatogonial proliferation through stem cell factor

**DOI:** 10.1101/2024.07.16.602181

**Authors:** Taichi Sugawara, Kayoko Sonoda, Nattapran Chompusri, Kazuhiro Noguchi, Seiji Okada, Mikio Furuse, Tomohiko Wakayama

## Abstract

Tight junctions (TJs) between adjacent Sertoli cells are believed to form immunological barriers that protect spermatogenic cells expressing autoantigens from autoimmune responses. However, there is no direct evidence that Sertoli cell TJs (SCTJs) do indeed form immunological barriers. Here, we analyzed male mice lacking claudin-11 (*Cldn11*), which encodes a SCTJ component, and found autoantibodies against antigens of spermatocytes/spermatids in their sera. Defective spermatogenesis was not restored in *Cldn11*-deficient mice on a genetic background mimicking a severely impaired adaptive immune system. This suggests that defective spermatogenesis is not caused by autoimmune responses against spermatogenic cells. Further analyses showed that *Cldn11* knockout impaired Sertoli cell polarization, localization of stem cell factor (SCF) (a key molecule for maintaining differentiating spermatogonia) to the basal compartment of seminiferous tubules, and also proliferation of differentiating spermatogonia. We propose that CLDN11 creates a microenvironment for SCF-mediated spermatogonial proliferation at the basal compartment via Sertoli cell polarization.

## Introduction

Cellular sheets of epithelia act as barriers separating the internal and external environments, generating distinct fluid compartments within the body. Tight junctions (TJs) are epithelial cell junctions formed at the most apical region of the lateral membrane. To establish compartmentalization, TJs restrict free diffusion of solutes through paracellular routes and thereby contribute to epithelial barrier function^1–4^. Electron microscopy using ultrathin sections shows that TJs are the region where two plasma membranes of neighboring cells fuse and obliterate the intercellular space^5^. While TJs are visualized as an anastomosing network of intramembranous particle strands (known as TJ strands) by freeze-fracture electron microscopy^6^. Claudins (CLDNs) are major integral membrane proteins constituting TJ strands^7,8^. CLDN1–8 have a PDZ-binding motif (YV sequence) in the C-terminus that binds to PDZ1 domains of the zonula occludens (ZO) family proteins (ZO1, ZO2, and ZO3), which are cytoplasmic scaffolding proteins^9^. Importantly, ZO1 and ZO2 are required not only for TJ formation but also the restriction of apical and basolateral membrane protein intermixing, which maintains epithelial polarity in cultured epithelial cells^10^. In addition to CLDNs, other integral membrane proteins such as occludin (OCLN) and junctional adhesion molecules (JAMs) are localized to TJs^11,12^.

In the testis, TJs form between adjacent Sertoli cells near the basement membrane of seminiferous tubules to establish the blood–testis barrier^13^. Sertoli cell TJs (SCTJs) are associated with ectoplasmic specializations composed of subsurface bundles of actin filaments and the more deeply located endoplasmic reticulum in the cytoplasm of Sertoli cells^13,14^. SCTJs physically divide seminiferous tubules into adluminal and basal compartments^13^. In the adluminal compartment, spermatocytes/spermatids express autoantigens on their cell surface^15,16^ and are sequestered from the systemic circulation by SCTJs. Thus, in the past several decades, SCTJs are believed to form an immunological barrier that protects spermatocytes/spermatids from the immune system by preventing autoimmune responses against spermatogenic cells. To unravel the role of SCTJs in spermatogenesis, several male mice lacking genes encoding TJ-associated transmembrane proteins have been generated. *Cldn11*-deficient mice exhibit smaller testes and defective spermatogenesis, leading to infertility^17,18^. Moreover, *Cldn3*-deficient mice have structurally and functionally intact SCTJs, and are capable of supporting spermatogenesis^19^. *Ocln*-deficient mice show atrophy in the seminiferous tubules when the mice reach around 40–60 weeks of age^20^. Deletion of *Jam1* results in subfertility due to defects in sperm tail formation and sperm motility, although the histological appearance of *Jam1*-deficient mouse testes is normal^21^. *Jam2*-deficient mice are fertile and exhibit normal spermatogenesis^22^. However, it remains unclear whether SCTJs act as an immunological barrier to support spermatogenesis.

Spermatogonia are divided into “undifferentiated” and “differentiating” populations in mice. Undifferentiated spermatogonia differentiate into preleptotene spermatocytes via differentiating spermatogonia in the basal compartment of seminiferous tubules. Preleptotene spermatocytes migrate into the adluminal compartment across SCTJs for further differentiation^23,24^. Stem cell factor (SCF) is a key molecule for preserving the population of differentiating spermatogonia^25^, and is encoded by the Steel (*Sl*) locus^26–28^. In particular, SCF from Sertoli cells, but not other somatic cells, including Leydig cells, endothelial cells, or smooth muscle cells, is indispensable for maintenance of differentiating spermatogonia^29^. Alternative splicing produces two *Scf* mRNAs, encoding either a transmembrane or secretory form^30^. In *Sl*/*Sl^d^* mutant mice, which express only the secretory form of SCF, development of male germ cells is impaired^30^, suggesting that a transmembrane form of SCF is critical for its biological function. Stable expression of a transmembrane form of SCF, in which GFP is inserted into the extracellular region, in cultured epithelial cells showed that this type of SCF localizes to the basolateral membrane^31^. However, it is largely unknown how SCF distributes on the Sertoli cell surface in seminiferous tubules and how its localization is regulated.

In the present study, we examined the role of CLDN11 in immunological barrier formation in the testis using *Cldn11*-deficient mice. We show that CLDN11 is necessary for SCTJ formation, Sertoli cell barrier function, and also prevents production of autoantibodies against antigens of spermatocytes/spermatids. These results support the idea that SCTJs restrict leakage of antigens from the adluminal compartment of seminiferous tubules to inhibit autoantibody production. Furthermore, we show that CLDN11 inhibits penetration of autoantibodies from the testicular interstitium into the seminiferous tubules, suggesting that CLDN11 acts as an immunological barrier to suppress the autoimmune response to spermatogenic cells. Importantly, however, our results demonstrate that CLDN11-mediated suppression of autoimmune responses are dispensable for spermatogenesis. Finally, we suggest that CLDN11 is required for Sertoli cell polarization and localization of SCF at the basal compartment of seminiferous tubules, contributing to proliferation of differentiating spermatogonia. Our findings have prompted us to reconsider the classical concept of SCTJ-mediated immunological barriers for spermatogenic cells, and shed new light on the role of CLDN11 in Sertoli cell polarization during spermatogenesis.

## Results

### CLDN11 is required for spermatogenesis in mice

CLDN11 is a component of SCTJs^17,32^. Published single-cell RNA sequencing (RNA-seq) data using spermatogenic cells from adult humans and adult mice suggest that human *CLDN11* and mouse *Cldn11* mRNA are predominantly expressed in Sertoli cells, but not spermatogenic cells of seminiferous tubules (Supplementary Fig. 1a–d)^33^. To identify the role of SCTJs in adult steady-state spermatogenesis, we used *Cldn11* knockout (*Cldn11*^-/-^) mice^34^. *Cldn11*^-/-^ mice had smaller sized testes and decreased testis weight compared with control mice (Fig. 1a). Immunohistochemical analyses using frozen testis sections showed loss of CLDN11 in seminiferous tubules of *Cldn11*^-/-^ mice (Supplementary Fig. 2a). We also confirmed complete loss of CLDN11 by western blotting using whole testis lysates prepared from *Cldn11*^-/-^ mice (Supplementary Fig. 2b). Histological examination revealed that *Cldn11* knockout impaired spermatogenesis in the testis and resulted in loss of spermatozoa in the cauda epididymis (Fig. 1b, c). These findings suggest that CLDN11 plays a pivotal role in spermatogenesis in mice. To further characterize this role, we performed immunohistochemistry using spermatogenic cell markers. We confirmed nearly complete loss of peanut agglutinin (PNA) lectin-positive cells (a marker of spermatids) in *Cldn11*^-/-^ mice (Supplementary Fig. 3a, b). We also observed a decrease in the number of VASA-positive cells (a marker of spermatids and spermatocytes) and synaptonemal complex protein 3 (SCP3)-positive cells (a marker of spermatocytes) in *Cldn11*^-/-^ mice (Supplementary Fig. 3c, d). In addition, the number of KIT-positive cells (a marker of differentiating spermatogonia) was significantly decreased in seminiferous tubules of *Cldn11*^-/-^ mice (Fig. 1d). The proportion of KIT-positive differentiating spermatogonia expressing Ki67, a proliferation marker, in *Cldn11*^-/-^ mice was comparable to *Cldn11*^+/-^ mice (Supplementary Fig. 4). This suggests that the proliferative capacity of residual KIT-positive differentiating spermatogonia was maintained in *Cldn11*^-/-^ mice. In contrast, the number of cells expressing undifferentiated spermatogonium markers, such as GDNF family receptor alpha 1 (GFRA1), promyelocytic leukemia zinc finger (PLZF), or Lin-28 homolog A (LIN28A), was largely unaffected in *Cldn11*^-/-^ mice (Fig. 1e and Supplementary Fig. 5a, b). To further investigate the cause for the decrease in spermatogenic cells in *Cldn11*^-/-^ mice, we performed terminal deoxynucleotidyl transferase-mediated dUTP nick-end labeling (TUNEL) assay to detect apoptotic cells. We found that *Cldn11*^-/-^ mice had a higher number of TUNEL-positive cells in seminiferous tubules compared with *Cldn11*^+/-^ mice (Fig. 1f). Furthermore, we detected TUNEL signals in VASA-positive and SCP3-positive cells of *Cldn11*^-/-^ mice, but not KIT-positive, PLZF-positive, or Wilms tumor 1 (WT1)-positive cells (a marker of Sertoli cells) (Fig. 1g and Supplementary Fig. 6). These results suggest that defective spermatogenesis in *Cldn11*^-/-^ mice is caused by a reduction in differentiating spermatogonia and an increase in apoptosis of spermatocytes. This demonstrates that CLDN11 is essential for adult steady-state spermatogenesis in mice.

**Fig. 1.**
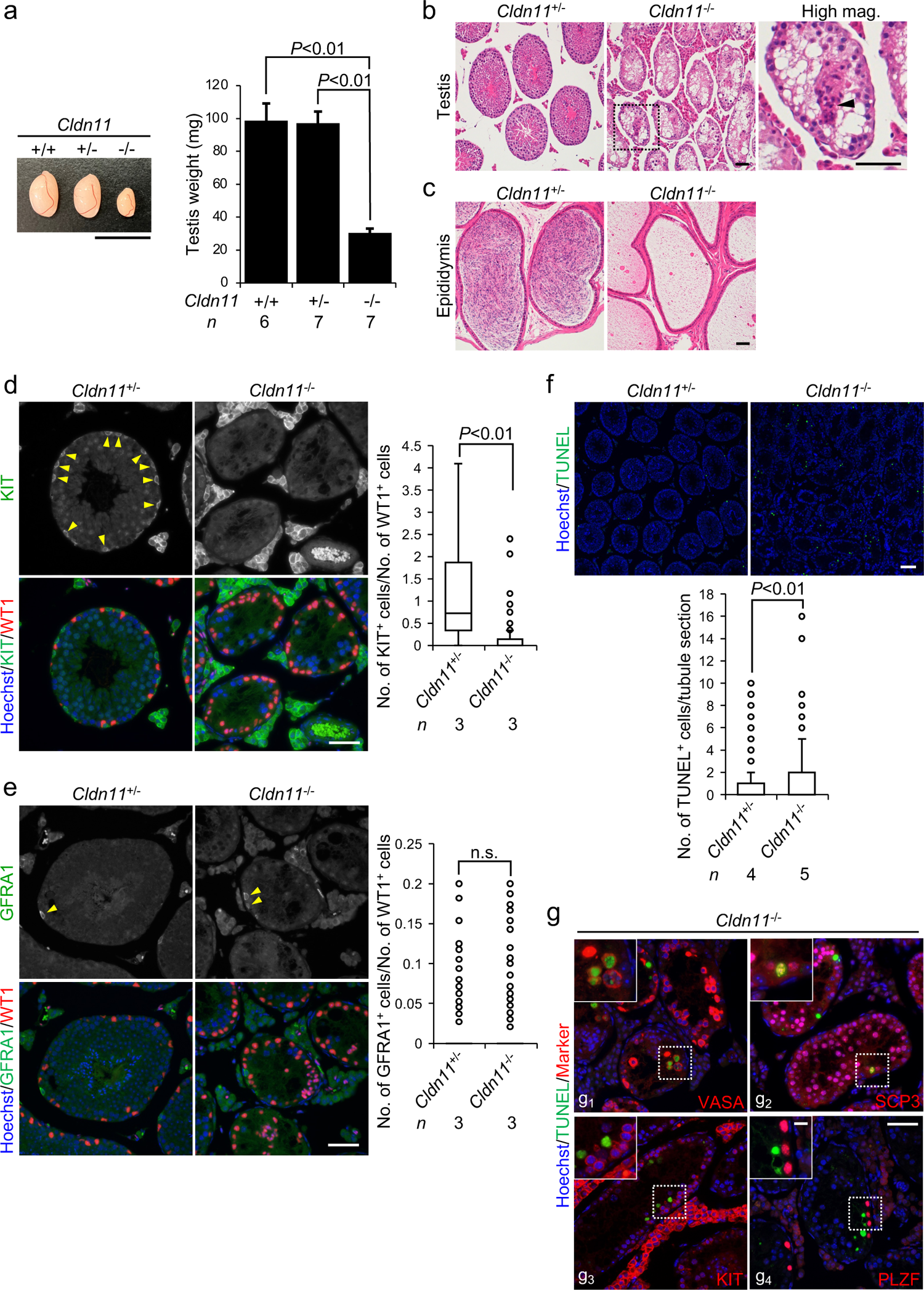
CLDN11 is required for spermatogenesis in mice. **a** Appearance and weight of testes from *Cldn11*^+/+^, *Cldn11*^+/-^, and *Cldn11*^-/-^ mice. The number of biologically independent mice in each group is shown as *n* in the graph. Data are shown as mean ± SD and were analyzed by Dunnett’s test. **b**, **c** Hematoxylin and eosin staining of sections prepared from testes (**b**) and the cauda epididymides (**c**) of *Cldn11*^+/-^ and *Cldn11*^-/-^ mice. The black dotted square is the region shown in a high magnification image (High mag.). A black arrowhead indicates a Sertoli cell cluster. **d**, **e** Immunohistochemistry of testis sections prepared from *Cldn11*^+/-^ and *Cldn11*^-/-^ mice using anti-KIT and anti-WT1 antibodies (**d**) or anti-GFRA1 and anti-WT1 antibodies (**e**). Cells expressing KIT (**d**) or GFRA1 (**e**) in seminiferous tubules are shown by yellow arrowheads. The number of KIT^+^ (**d**) or GFRA1^+^ (**e**) cells on seminiferous tubule sections from testes of *Cldn11*^+/-^ and *Cldn11*^-/-^ mice were normalized to the number of WT1^+^ Sertoli cells in the graph. A total of 228–247 seminiferous tubules on testis sections from three biologically independent *Cldn11*^+/-^ or *Cldn11*^-/-^ mice were analyzed. Data were analyzed by Mann–Whitney *U* test. n.s. (not significant): *P* > 0.05. (**f**) TUNEL assay using testis sections prepared from *Cldn11*^+/-^ and *Cldn11*^-/-^ mice. The number of TUNEL^+^ cells per seminiferous tubule section was counted. A total of 541 or 709 seminiferous tubules on testis sections from four *Cldn11*^+/-^ or five *Cldn11*^-/-^ biologically independent mice, respectively, were analyzed. Data were analyzed by Mann–Whitney *U* test. (**g**) TUNEL assay with immunohistochemistry of testis sections from *Cldn11*^-/-^ mice using anti-VASA (**g_1_**), anti-SCP3 (**g_2_**), anti-KIT (**g_3_**), or anti-PLZF antibodies (**g_4_**). Insets are magnified images of the regions indicated by white dotted squares. Scale bars: 1 cm (**a**), 50 μm (**b**– **e**, **g**), 100 μm (**f**), and 10 μm (**g**, inset).

Next, we examined the role of CLDN11 during the first wave of spermatogenesis. We investigated localization of CLDN11 and ZO1 in seminiferous tubules of postnatal day (P)-10, P20, and P30 wild-type mice by immunohistochemistry. At P10, CLDN11 and ZO1 signals were detected within seminiferous tubules but did not accumulate near the basement membrane of tubules (Supplementary Fig. 7a). However, at P20 and P30, CLDN11 colocalized with ZO1 near the basement membrane (Supplementary Fig. 7a), suggesting that SCTJs form between P10 and P20 in mice. Our observations are consistent with previous results obtained by electron microscopy analyses of mouse testes^35,36^. Although testis weight was not significantly altered at P10, P20, or P30 in *Cldn11*^-/-^ mice compared with control mice (Supplementary Fig. 7b), histological abnormalities, including Sertoli cell clusters at the adluminal compartment of seminiferous tubules, were found at P20 and P30, but not P10, in *Cldn11*^-/-^ mice (Supplementary Fig. 7c). No elongated spermatids were observed at P30 in *Cldn11*^-/-^ mice, but were present at P30 in *Cldn11*^+/-^ mice (Supplementary Fig. 7c). Collectively, our findings suggest that CLDN11-mediated SCTJ formation is necessary to accomplish the first wave of spermatogenesis, although CLDN11 is dispensable for its onset.

### CLDN11 is required for SCTJ formation and Sertoli cell barrier function

To clarify the effect of *Cldn11* knockout on SCTJ formation, we examined localization of TJ-associated proteins in seminiferous tubules. Immunohistochemical analyses of frozen testis sections showed that OCLN and ZO1 still colocalized near the basement membrane of seminiferous tubules in *Cldn11*^-/-^ mice (Supplementary Fig. 8a). Colocalization of JAM1 and ZO1 decreased with *Cldn11* knockout (Supplementary Fig. 8b). In addition to CLDN11, CLDN3 and CLDN5 are thought to be expressed in mouse Sertoli cells^37,38^. Thus, we investigated localization of CLDN3 and CLDN5. We found partial colocalization of CLDN3 with ZO1 in seminiferous tubules of wild-type mice, which was lost in *Cldn11*^-/-^ mice (Supplementary Fig. 8c). Prominent CLDN5 signals were not detected in seminiferous tubules of wild-type or *Cldn11*^-/-^ mice (Supplementary Fig. 8d). Next, we examined ZO1 localization by whole-mount immunofluorescence of seminiferous tubules. ZO1 showed junctional localization in seminiferous tubules of *Cldn11*^+/-^ mice. Intriguingly, however, ZO1 localization was discontinuous and fragmented in seminiferous tubules of *Cldn11*^-/-^ mice (Fig. 2a), raising the possibility that CLDN11 regulates ZO1 localization. To examine this, we overexpressed mouse CLDN11 in mouse L fibroblasts lacking cadherin-mediated cell adhesion^39^. In L fibroblasts overexpressing CLDN11, ZO1 was recruited to cell–cell contacts and colocalized with CLDN11 (Fig. 2b). To determine whether CLDN11 binds to ZO1, similar to other CLDNs (CLDN1–8) with YV sequences in their C-termini^9^, we next performed *in vitro* binding assays. We generated bacterial glutathione S-transferase (GST) fusion proteins for the C-terminal 29 amino acids of mouse CLDN11 (aa 179–207: GST-CL11) and its deletion mutant lacking the two C-terminal amino acids corresponding to a putative PDZ-binding motif (HV sequence) (aa 179–205: GST-CL11ΔHV). To determine which PDZ domain of ZO1 binds to CLDN11, we also generated bacterial maltose-binding protein (MBP) fusion proteins for PDZ1 (aa 19–113: MBP-zPDZ1), PDZ2 (aa 181–292: MBP-zPDZ2), and PDZ3 (aa 423–503: MBP-zPDZ3) of ZO1 (Fig. 2c). GST pull-down assays showed that GST-CL11 bound to MBP-zPDZ1 (Fig. 2d), while GST-CL11ΔHV did not (Fig. 2e), indicating that CLDN11 directly binds to PDZ1 of ZO1 via its PDZ-binding motif (HV sequence). These findings support the idea that CLDN11 directly regulates ZO1 localization in seminiferous tubules.

**Fig. 2.**
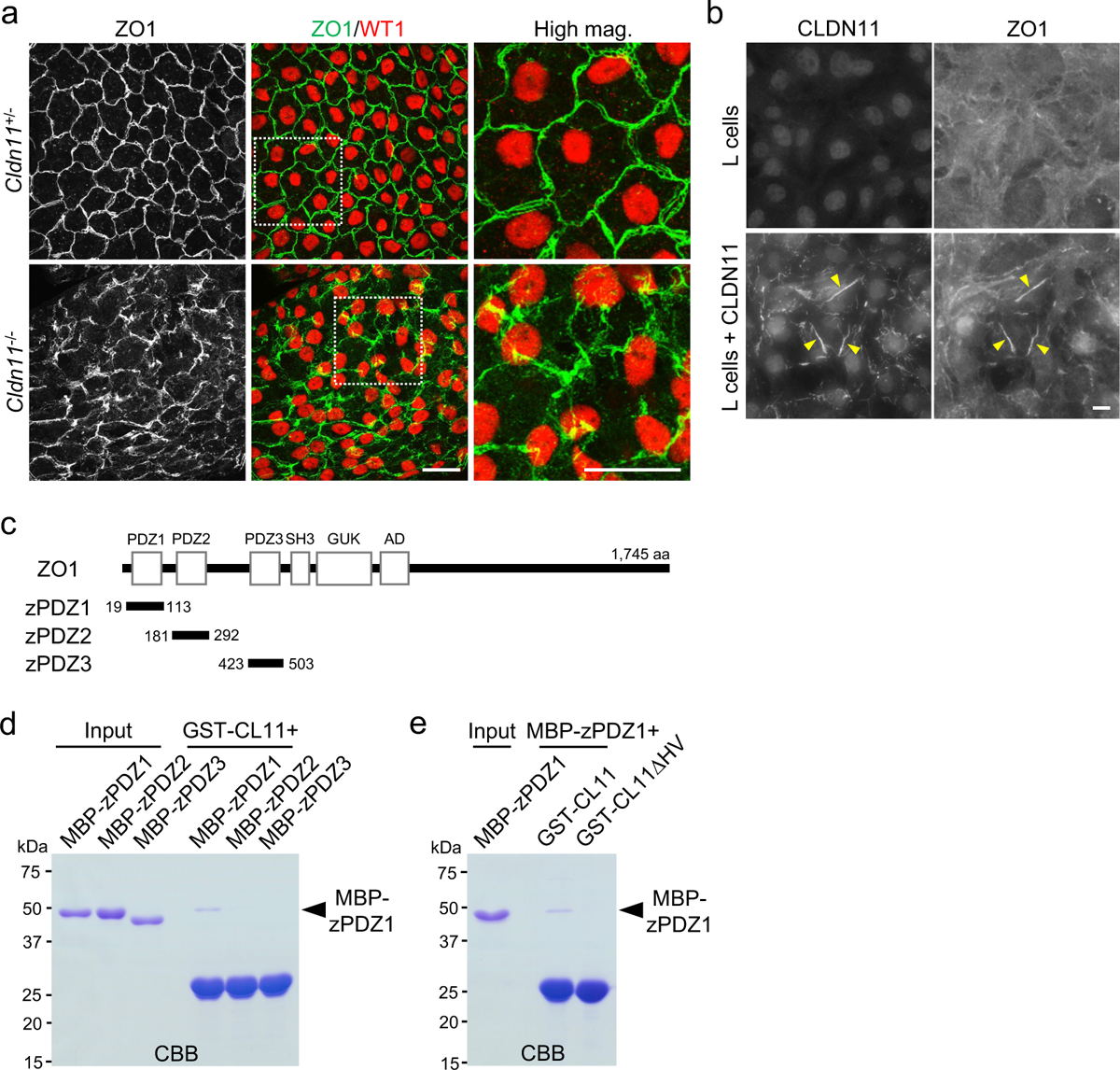
CLDN11 regulates ZO1 localization in seminiferous tubules. **a** Whole-mount immunofluorescence staining of seminiferous tubules from *Cldn11*^+/-^ and *Cldn11*^-/-^ mice using anti-ZO1 and anti-WT1 antibodies. White dotted squares indicate the regions shown in high magnification images (High mag.). **b** Immunofluorescence staining of parental L cells and L cells stably expressing mouse CLDN11 using anti-CLDN11 and anti-ZO1 antibodies. Yellow arrowheads show colocalization of CLDN11 with ZO1. **c** Schematic diagram of the domain structure of the mouse ZO1 protein (1,745 amino acids). The N-terminal portion of ZO1 contains three PDZ domains, an SH3 domain, a guanylate kinase (GUK) domain, and an acidic domain (AD). Distinct portions of the ZO1 protein were fused to maltose-binding protein (MBP); shown by three black lines with amino acid numbers. **d**, **e** Glutathione S-transferase (GST)-fused proteins of aa 179–207 and aa 179–205 of mouse CLDN11 were designated as GST-CL11 and GST-CL11ΔHV, respectively. MBP-fused proteins of PDZ1 (aa 19–113), PDZ2 (aa 181–292), and PDZ3 domain (aa 423–503) of ZO1 were designated as MBP-zPDZ1, MBP-zPDZ2, and MBP-zPDZ3, respectively. Input: purified MBP-fused proteins. GST pull-down assays were performed using GST-CL11, MBP-zPDZ1, MBP-zPDZ2, and MBP-zPDZ3 (**d**) or GST-CL11, GST-CL11ΔHV, and MBP-zPDZ1 (**e**). The resulting samples were separated by SDS–PAGE followed by Coomassie Brilliant Blue (CBB) staining. Scale bars: 30 μm (**a**) and 10 μm (**b**).

To investigate the effect of *Cldn11* knockout on the ultrastructure of SCTJs, we used ultrathin sections for electron microscopy of seminiferous tubules. In *Cldn11*^+/-^ mice, close contacts between neighboring plasma membranes of Sertoli cells were observed (Fig. 3a). However, these plasma membrane contacts were lost in *Cldn11*^-/-^ mice, with large gaps observed between Sertoli cell plasma membranes associated with ectoplasmic specializations (Fig. 3a). Next, we analyzed Sertoli cell barrier function by injection of a tracer, specifically a primary amine-reactive biotinylation reagent (557 Da)^40^, into the interstitium of the testis. Diffusion of the biotinylation reagent from the interstitium stopped at ZO1-positive SCTJs in *Cldn11*^+/-^ mice (Fig. 3b). In contrast, the biotinylation reagent reached the adluminal compartment of seminiferous tubules in *Cldn11*^-/-^ mice (Fig. 3c). Taken together, these findings indicate that CLDN11 is essential for SCTJ formation and Sertoli cell barrier function.

**Fig. 3.**
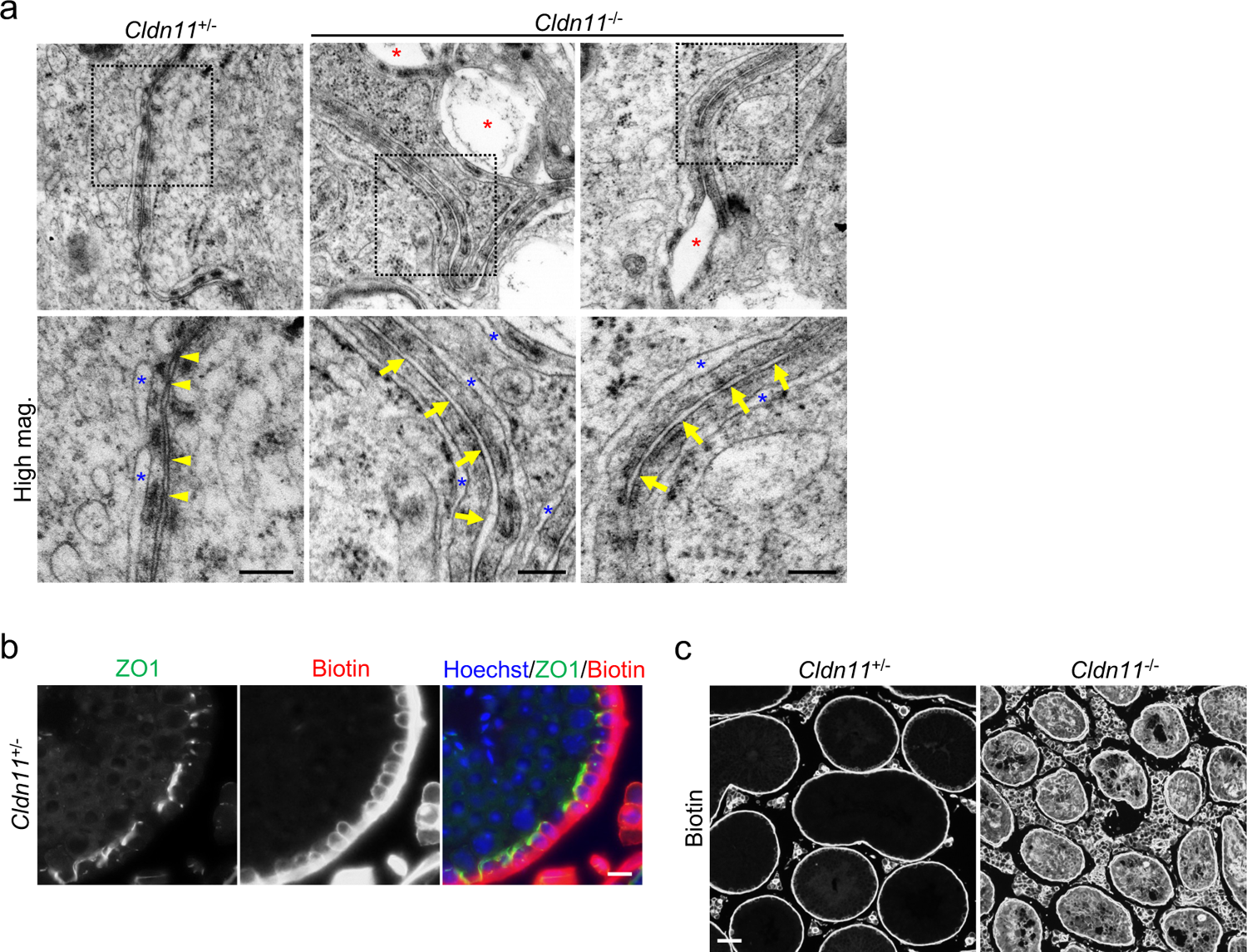
CLDN11 is required for SCTJ formation and Sertoli cell barrier function. **a** Ultrathin section electron microscopy of seminiferous tubules from *Cldn11*^+/-^ and *Cldn11*^-/-^ mice. Yellow arrowheads indicate close contacts between neighboring plasma membranes of Sertoli cells. Yellow arrows indicate intercellular spaces without plasma membrane contacts between adjacent Sertoli cells. Blue asterisks indicate endoplasmic reticulum cisternae constituting ectoplasmic specializations. Red asterisks indicate large gaps between Sertoli cell plasma membranes associated with ectoplasmic specializations. **b**, **c** Biotin tracer assay. Sulfo-NHS-LC-Biotin was injected into the interstitium of testes from *Cldn11*^+/-^ and *Cldn11*^-/-^ mice. Immunohistochemistry of testis sections from *Cldn11*^+/-^ mice using anti-ZO1 antibody and streptavidin conjugated to a fluorescent dye (**b**). Testis sections from *Cldn11*^+/-^ and *Cldn11*^-/-^ mice were labeled with streptavidin (**c**). Scale bars: 200 nm (**a**), 10 μm (**b**), and 50 μm (**c**).

### CLDN11 prevents production of autoantibodies against antigens in spermatocytes/spermatids and penetration of IgG into seminiferous tubules

SCTJs are thought to act as an immunological barrier by restricting leakage of autoantigens in spermatocytes/spermatids from the adluminal compartment of seminiferous tubules. Thus, structural and functional defects in SCTJs are expected to generate autoantibodies against autoantigens. To examine this, we investigated whether autoantibodies are present in sera from *Cldn11*^-/-^ mice by immunohistochemistry of wild-type testis sections. Antigens were not detected within seminiferous tubules in 24 of 26 sera from control mice (Fig. 4a). In contrast, 3 of 20 sera and 4 of 20 sera from *Cldn11*^-/-^ mice reacted to antigens in nuclei of SCP3-positive cells or at the acrosome labeled with PNA lectin, respectively (Fig. 4b). In addition, we detected dot-like signals or signals from flagella of elongated spermatids at the adluminal compartment of wild-type seminiferous tubules using 2 of 20 sera from *Cldn11*^-/-^ mice as primary antibodies (Supplementary Fig. 9a). We also observed fluorescence signals similar to that in Fig. 4b_1_ or Supplementary Fig. 9a_2_ by immunohistochemistry using 2 of 26 sera from control mice (data not shown). Considering that autoantibodies against antigens of spermatocytes/spermatids are present in 9 of 20 sera from *Cldn11*^-/-^ mice compared with 2 of 26 sera from control mice (*P* < 0.01, Fisher’s exact test), our findings suggest that *Cldn11*^-/-^ mice tend to produce autoantibodies. Moreover, western blotting of whole wild-type testis lysates using sera from *Cldn11*^-/-^ mice detected extra bands that were not present using sera from *Cldn11*^+/-^ mice (Fig. 4c). This suggests the presence of autoantibodies against testicular antigens in sera from *Cldn11*^-/-^ mice. Next, we tested the possibility that autoantibodies against antigens of spermatocytes/spermatids penetrate into the adluminal compartment of seminiferous tubules in *Cldn11*^-/-^ mice. Since it is difficult to visualize autoantibodies themselves, we investigated distribution of IgG by immunohistochemistry of frozen testis sections. We found that IgG localized outside CLDN11-positive SCTJs in *Cldn11*^+/-^ mice (Fig. 4d), while IgG accumulated at the adluminal compartment of seminiferous tubules in *Cldn11*^-/-^ mice (Fig. 4e). In addition, tracer assays showed that flux of 150 kDa dextran, which has about the same molecular weight as IgG, from the interstitium into the adluminal compartment of seminiferous tubules was increased in *Cldn11*^-/-^ mouse testes (Fig. 4f). These results suggest that CLDN11 forms a physical barrier to macromolecules such as IgG, including autoantibodies. Meanwhile, CD45-positive cells (a marker of leukocytes) were not observed in seminiferous tubules of *Cldn11*^+/-^ or *Cldn11*^-/-^ mice (Supplementary Fig. 9b), suggesting that CLDN11 is dispensable for preventing infiltration of leukocytes into seminiferous tubules.

**Fig. 4.**
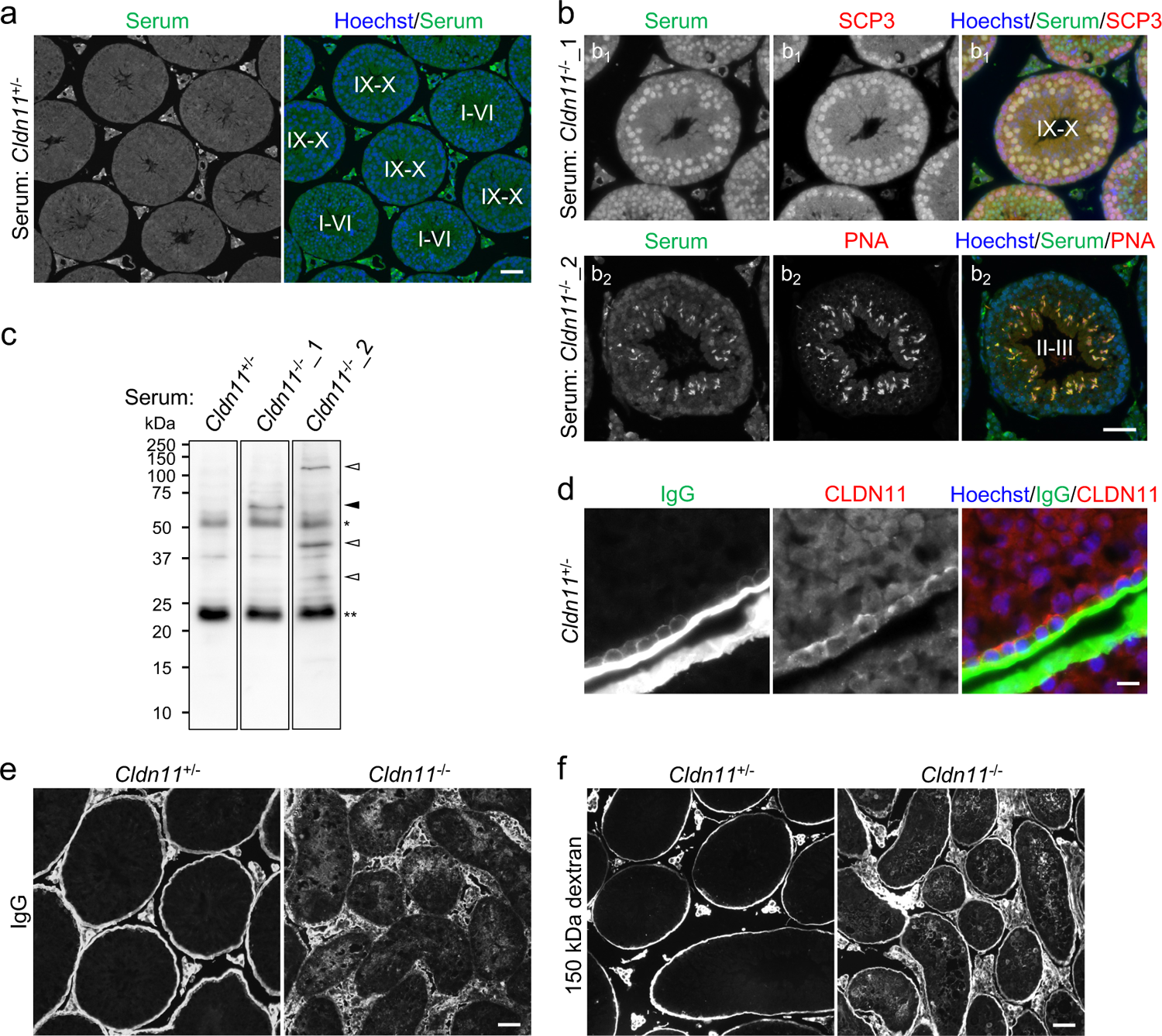
Production of autoantibodies against antigens of spermatocytes/spermatids and penetration of IgG into seminiferous tubules in *Cldn11*^-/-^ mice. **a** Immunohistochemistry of wild-type testis sections using serum from a *Cldn11*^+/-^ mouse as a primary antibody. Stages of spermatogenesis are shown by Roman numerals. **b** Immunohistochemistry of wild-type testis sections using serum from a *Cldn11*^-/-^ mouse (*Cldn11*^-/-^_1) and anti-SCP3 antibody (**b_1_**), or serum from a *Cldn11*^-/-^ mouse (*Cldn11*^-/-^_2) and PNA lectin (**b_2_**). Stages of spermatogenesis are shown by Roman numerals. **c** Western blotting of wild-type testis lysates with serum from a *Cldn11*^+/-^ mouse and sera from *Cldn11*^-/-^ mice used in (**b**). A black arrowhead and white arrowheads indicate testicular antigens detected by sera in (**b_1_**) and (**b_2_**), respectively. *IgG heavy chains. **IgG light chains. **d** Immunohistochemistry of frozen testis sections from *Cldn11*^+/-^ mice using anti-mouse IgG and anti-CLDN11 antibodies. **e** Immunohistochemistry of frozen testis sections from *Cldn11*^+/-^ and *Cldn11*^-/-^ mice using anti-mouse IgG antibody. **f** Tracer assay. Dextran (150 kDa) conjugated to a fluorescent dye was injected into the interstitium of testes from *Cldn11*^+/-^ and *Cldn11*^-/-^ mice. Scale bars: 50 μm (**a**, **b**, **e**, **f**) and 10 μm (**d**).

### *Rag2* knockout does not restore defective spermatogenesis caused by *Cldn11* knockout

Our findings suggest that CLDN11 prevents production of autoantibodies against antigens of spermatocytes/spermatids, which penetrate into the adluminal compartment of seminiferous tubules. This raises the possibility that CLDN11 contributes to spermatogenesis by inhibiting autoimmune responses against spermatogenic cells with autoantigens. To test this, we generated *Cldn11*^-/-^/recombination activating gene 2 (*Rag2*)^-/-^ male mice by crossing *Cldn11*^-/-^ mice and *Rag2*^-/-^/Janus kinase 3 (*Jak3*)^-/-^ mice^41^, in which spermatogenesis occurs normally (Supplementary Fig. 10a–c). *Rag2*^-/-^ mice fail to produce mature T and B lymphocytes due to an inability to initiate V(D)J rearrangement, leading to a severe combined immune deficient phenotype^42^. Thus, on a *Rag2* knockout background, *Cldn11*^-/-^ mice are expected to lack an adaptive immune response. Testis size was smaller in *Cldn11*^-/-^ mice and *Cldn11*^-/-^/*Rag2*^-/-^ mice compared with control mice (Fig. 5a). There was also no significant difference in testis weight between *Cldn11*^-/-^ mice and *Cldn11*^-/-^/*Rag2*^-/-^ mice (Fig. 5a). As expected, western blotting of whole testis lysates showed that IgG proteins were completely lost on a *Rag2* knockout background (Fig. 5b). *Cldn11*^-/-^/*Rag2*^-/-^ mice exhibited abnormal testis histology and complete loss of spermatozoa in the cauda epididymides, similar to *Cldn11*^-/-^ mice (Fig. 5c, d). In addition, there were no notable differences in populations of PNA lectin-positive cells and SCP3-positive cells between *Cldn11*^-/-^ mice and *Cldn11*^-/-^/*Rag2*^-/-^ mice (Fig. 5e). These results demonstrate that CLDN11-mediated spermatogenesis is independent of RAG2 function, and also suggest that humoral and cell-mediated autoimmune responses against spermatogenic cells with autoantigens are not a cause of defective spermatogenesis in *Cldn11*^-/-^ mice.

**Fig. 5.**
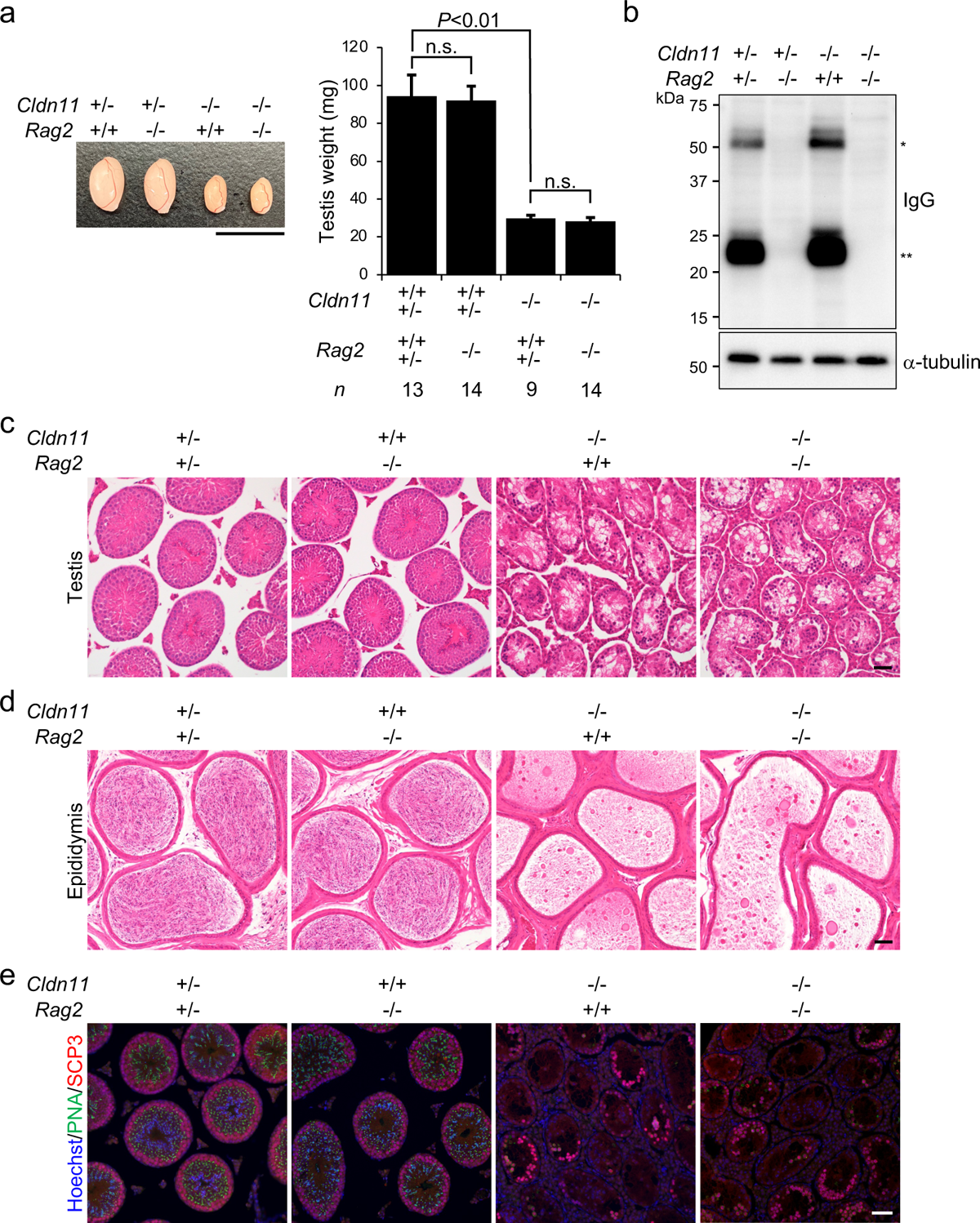
Defective spermatogenesis in *Cldn11*^-/-^ mice is not restored by additional *Rag2* knockout. **a** Appearance of testes from *Cldn11*^+/-^/*Rag2*^+/+^, *Cldn11*^+/-^/*Rag2*^-/-^, *Cldn11*^-/-^/*Rag2*^+/+^, and *Cldn11*^-/-^/*Rag2*^-/-^ mice. Testis weight of mice from each genotype is shown in the graph. The number of biologically independent mice in each group is shown as *n* in the graph. Data are shown as mean ± SD and were analyzed by Tukey–Kramer test. n.s. (not significant): *P* > 0.05. **b** Western blotting of testis lysates prepared from *Cldn11*^+/-^/*Rag2*^+/-^, *Cldn11*^+/-^/*Rag2*^-/-^, *Cldn11*^-/-^/*Rag2*^+/+^, and *Cldn11*^-/-^/*Rag2*^-/-^ mice using anti-mouse IgG and anti-α-tubulin antibodies. *IgG heavy chains. **IgG light chains. **c**, **d** Hematoxylin and eosin staining of sections prepared from testes (**c**) and the cauda epididymides (**d**) of *Cldn11*^+/-^/*Rag2*^+/-^, *Cldn11*^+/+^/*Rag2*^-/-^, *Cldn11*^-/-^/*Rag2*^+/+^, and *Cldn11*^-/-^/*Rag2*^-/-^ mice. **e** Immunohistochemistry of testis sections from *Cldn11*^+/-^/*Rag2*^+/-^, *Cldn11*^+/+^/*Rag2*^-/-^, *Cldn11*^-/-^/*Rag2*^+/+^, and *Cldn11*^-/-^/*Rag2*^-/-^ mice using PNA lectin and anti-SCP3 antibody. Scale bars: 1 cm (**a**) and 50 μm (**c**, **d**, **e**).

### CLDN11 regulates Sertoli cell polarization and localization of cell junctional components in Sertoli cells

Analyses of *ZO1*/*ZO2* double knockout Madin–Darby canine kidney (MDCK) II epithelial cells previously showed that ZO1/ZO2 are required for epithelial polarization, and can prevent apical and basolateral membrane protein intermixing on the cell surface^10^. Since *Cldn11* knockout severely impairs junctional localization of ZO1 in seminiferous tubules (Fig. 2a), we examined the role of CLDN11 in Sertoli cell polarization. To clarify the precise localization of membrane proteins in Sertoli cells, we depleted spermatogenic cells in the testis of *Cldn11*^+/-^ and *Cldn11*^-/-^ mice by busulfan treatment. Immunohistochemical analyses showed that ezrin (EZR), an apical marker, was located at the adluminal compartment in seminiferous tubules of busulfan-treated *Cldn11*^+/-^ mice (Fig. 6a). However, EZR mislocalized near the basement membrane in seminiferous tubules of busulfan-treated *Cldn11*^-/-^ mice (Fig. 6a). The basolateral marker, Na/K ATPase α1 subunit (ATP1A1), localized to the basolateral membrane of Sertoli cells, with some ATP1A1 closely localized with a TJ marker in seminiferous tubules of busulfan-treated *Cldn11*^+/-^ mice (Fig. 6b). In contrast, ATP1A1 diffusely localized to adluminal and basal regions in seminiferous tubules of busulfan-treated *Cldn11*^-/-^ mice (Fig. 6b). A polarity protein complex composed of PAR3/PAR6/aPKC localizes to TJs^43^. However, we could not confirm SCTJ localization of polarity proteins by immunohistochemistry of wild-type testis sections (data not shown). The mRNA expression levels of *Par3*, *Par6a*, *Prkcz*, and *Prkci* are quite low in Sertoli cells, as indicated by published single-cell RNA-seq data using spermatogenic cells from adult mice (Supplementary Fig. 11)^33^, which may explain this discrepancy. We also found that *Cldn11* knockout affected localization of nectin-2 (NECTIN2), an adherens junction protein, and gap junction α1 protein (GJA1), a gap junction protein, in seminiferous tubules. In busulfan-treated *Cldn11*^+/-^ mice, NECTIN2 and GJA1 closely localized with a TJ marker near the basement membrane in seminiferous tubules (Fig. 6c, d). However, in busulfan-treated *Cldn11*^-/-^ mice, NECTIN2 diffusely localized to the adluminal and basal regions (Fig. 6c), while GJA1 mislocalized from Sertoli cell–Sertoli cell junctions (Fig. 6d). These results suggest that CLDN11 regulates Sertoli cell polarization and localization of cell junctional components in Sertoli cells.

**Fig. 6.**
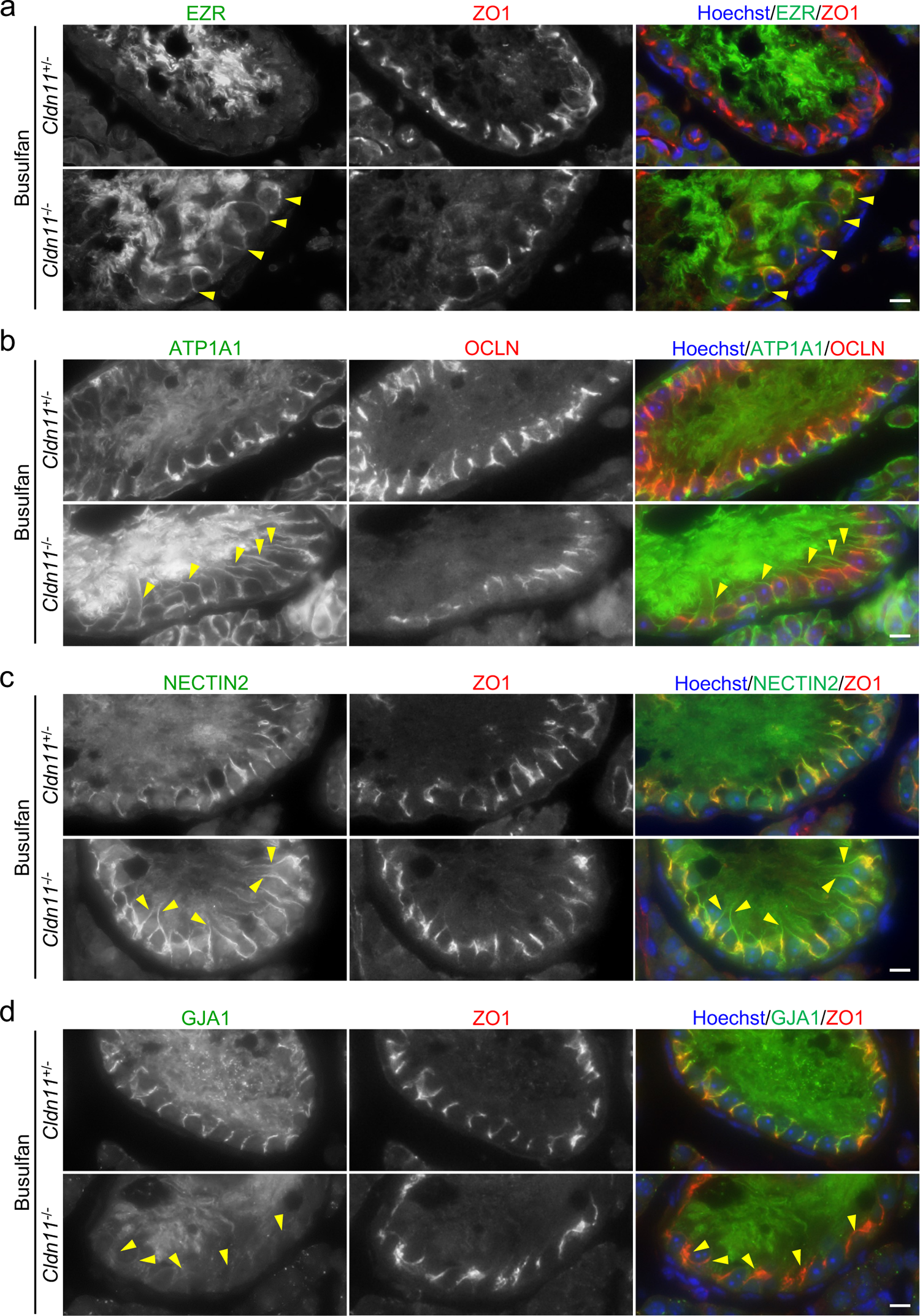
Epithelial polarity markers and components of junctional complexes are mislocalized in Sertoli cells of busulfan-treated *Cldn11*^-/-^ mice. **a**–**d** Immunohistochemistry of testis sections prepared from busulfan-treated *Cldn11*^+/-^ and *Cldn11*^-/-^ mice using anti-EZR and anti-ZO1 (**a**), anti-ATP1A1 and anti-OCLN (**b**), anti-NECTIN2 and anti-ZO1 (**c**), or anti-GJA1 and anti-ZO1 antibodies (**d**). Yellow arrowheads show mislocalization of EZR near the basement membrane (**a**), diffuse localization of ATP1A1 (**b**) and NECTIN2 (**c**) to the adluminal region, and delocalization of GJA1 from Sertoli cell–Sertoli cell junctions (**d**) in seminiferous tubules of busulfan-treated *Cldn11*^-/-^ mice. Scale bars: 10 μm (**a**–**d**).

### CLDN11 regulates proliferation of differentiating spermatogonia through SCF

It has been suggested that SCF, a ligand of the KIT receptor, is necessary for proliferation of differentiating spermatogonia, but not undifferentiated spermatogonia^25^. Taking into consideration that the number of differentiating spermatogonia was reduced in *Cldn11*^-/-^ mice (Fig. 1d), we hypothesized that CLDN11 maintains differentiating spermatogonia through SCF. First, we attempted to confirm the role of SCF in spermatogenesis using *Sl*/*Sl^d^*mutant mice lacking a transmembrane form of SCF^30^. Immunohistochemistry showed that PLZF-positive undifferentiated spermatogonia resided in some seminiferous tubules on testis sections prepared from *Sl*/*Sl^d^* mutant mice (Supplementary Fig. 12a, b), consistent with previous observations^25^. In the seminiferous tubules, LIN28A-positive undifferentiated spermatogonia were also observed (Supplementary Fig. 12c), while a population of KIT-positive differentiating spermatogonia was decreased (Supplementary Fig. 12d). This suggests that a transmembrane form of SCF is necessary for maintenance of differentiating spermatogonia. A recent study demonstrated that SCF in Sertoli cells, but not other somatic cells in the testis, is required for maintenance of differentiating spermatogonia^29^. Thus, we examined SCF localization in Sertoli cells of busulfan-treated testes. Immunohistochemical analyses showed that SCF closely localized with a TJ marker near the basement membrane in busulfan-treated *Cldn11*^+/-^ testes (Fig. 7a). In contrast, SCF mislocalized from Sertoli cell–Sertoli cell junctions in busulfan-treated *Cldn11*^-/-^ testes (Fig. 7a). To further clarify SCF localization in polarized epithelial cells, we generated MDCK II cells stably expressing a transmembrane form of SCF tagged with C-terminal hemagglutinin (HA) (Supplementary Fig. 13a). Immunofluorescence showed that HA-tagged SCF localized to the lateral membrane in MDCK II cells (Supplementary Fig. 13b, c). These results suggest that SCF preferentially localizes at the basal compartment of seminiferous tubules, and its localization is regulated by CLDN11. Next, to test the possibility that a reduction in the number of differentiating spermatogonia in *Cldn11*^-/-^ mice is caused by SCF delocalization from the basal compartment, we treated seminiferous tubules of *Cldn11*^-/-^ mice with recombinant SCF proteins corresponding to the extracellular region. Recombinant protein was administered from the interstitium for 10 days using gel beads labeled with a fluorescent dye^44^. To distinguish areas of seminiferous tubules close to bovine serum albumin (BSA)- or SCF-soaked beads, the areas were labeled with a red fluorescent dye, CM-DiI (Fig. 7b). The cell density of KIT-positive differentiating spermatogonia in seminiferous tubules labeled with CM-DiI was significantly increased by transplantation of SCF-soaked beads, but not BSA-soaked beads (Fig. 7c, d). Taken together, our findings suggest that CLDN11 regulates proliferation of differentiating spermatogonia through SCF (Fig. 7e).

**Fig. 7.**
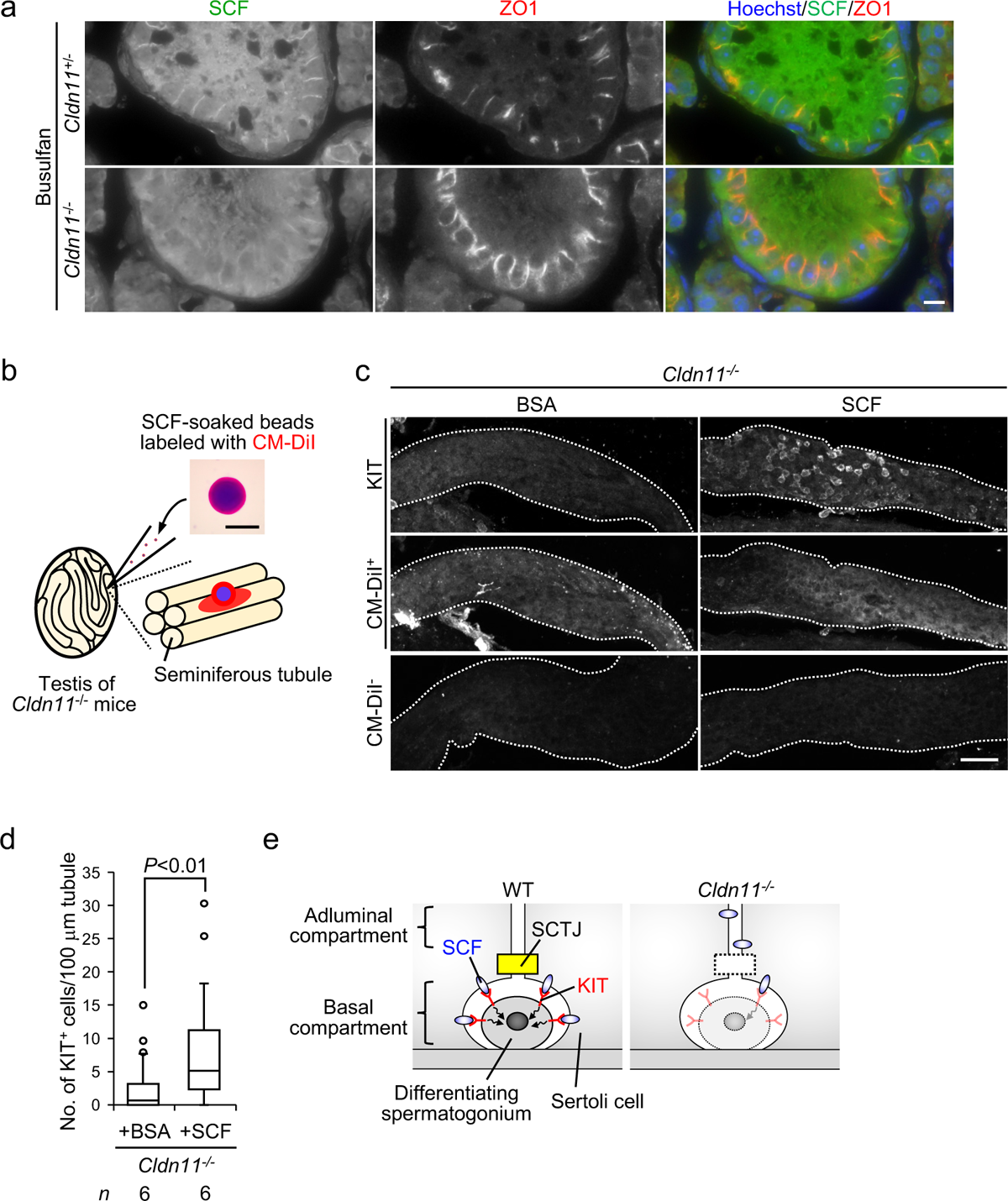
*In vivo* transplantation of SCF-soaked/CM-DiI-labeled beads into the testicular interstitium induces proliferation of differentiating spermatogonia in *Cldn11*^-/-^ mice. **a** Immunohistochemistry of testis sections prepared from busulfan-treated *Cldn11*^+/-^ and *Cldn11*^-/-^ mice using anti-SCF and anti-ZO1 antibodies. **b** Schema for *in vivo* transplantation of SCF-soaked/CM-DiI-labeled beads into the interstitium of *Cldn11*^-/-^ mouse testes. Appearance of a SCF-soaked/CM-DiI-labeled bead is shown. CM-DiI labels seminiferous tubules in proximity to the beads. **c** Whole-mount immunofluorescence staining using anti-KIT antibody on seminiferous tubules from *Cldn11*^-/-^ mouse testes, with bovine serum albumin (BSA)- or SCF-soaked beads transplanted into the interstitium. Seminiferous tubules with or without CM-DiI labeling are shown as CM-DiI^+^ and CM-DiI^-^, respectively. White dotted lines outline the seminiferous tubules. **d** BSA- and SCF-soaked beads were separately transplanted into distinct testes of a *Cldn11*^-/-^ mouse. Transplantation was performed using six biologically independent *Cldn11*^-/-^ mice. After, 67 and 72 seminiferous tubules labeled with CM-DiI derived from BSA- or SCF-soaked beads, respectively, were analyzed. The number of KIT^+^ cells per 100 μm seminiferous tubule labeled with CM-DiI was quantified. Data were analyzed by Mann–Whitney *U* test. **e** Working model showing the role of CLDN11 in maintenance of differentiating spermatogonia. CLDN11-based Sertoli cell tight junctions (SCTJs) physically divide seminiferous tubules into adluminal and basal compartments. KIT-positive differentiating spermatogonia are located at the basal compartment. Interaction of Sertoli cell-derived SCF with KIT is essential for survival of differentiating spermatogonia. SCF accumulates at the basal compartment through Sertoli cell polarization after SCTJ formation, presumably promoting KIT activation in differentiating spermatogonia. *Cldn11* knockout delocalizes SCF from the basal compartment and reduces the number of differentiating spermatogonia, likely due to a decrease in KIT activity. Scale bars: 10 μm (**a**), 100 μm (**b**), and 50 μm (**c**).

## Discussion

Classically, SCTJs have been thought to form immunological barriers to sequester autoantigens of spermatocytes/spermatids within the adluminal compartment, leading to prevention of autoimmune responses against spermatogenic cells. This well-known concept was established based upon certain observations. First, SCTJs are appropriate structures to restrict the permeability of autoantigens, autoantibodies, and immune cells through the paracellular pathway by eliminating the intercellular space. Second, SCTJ formation and the appearance of pachytene spermatocytes occurs around the same time (P14–P16) in seminiferous tubules of mice^35,36,45^, implying that SCTJs protect spermatocytes from the immune system. However, previous studies using mice lacking TJ-associated genes, such as *Cldn11*, *Cldn3*, *Ocln*, *Jam1*, or *Jam2*, have not provided direct evidence of the immunological barrier function of SCTJs^17–22^. In the present study, we show that *Cldn11* knockout induces production of autoantibodies against antigens expressed in spermatocytes/spermatids, suggesting that CLDN11 acts as an immunological barrier and likely restricts leakage of autoantigens from the adluminal compartment of seminiferous tubules into the systemic circulation. Moreover, previous studies show the likely involvement of SCTJs in immunological barrier formation. Analyses of mice with Sertoli cell-specific ablation of the androgen receptor demonstrate that the receptor in Sertoli cells regulates SCTJ formation at the ultrastructural level and inhibit production of autoantibodies against germ cell antigens^46^. This suggests that androgen receptor-mediated maintenance of testicular immune privilege may depend on SCTJ structures. Furthermore, mice with conditional knockout of rho guanine nucleotide exchange factor 15 (*Arhgef15*) in Sertoli cells exhibit disruption of Sertoli cell barrier function and defects in testicular immune homeostasis characterized by production of autoantibodies against testicular antigens^47^. These results imply that ARHGEF15 may maintain immune homeostasis in the testis through SCTJ function. In addition to SCTJs, it is suggested that regulatory T cells (Tregs) play an important role in suppression of the production of autoantibodies against spermatogenic cell antigens^48,49^. Considering that Sertoli cells can induce Treg conversion and expansion in a transforming growth factor (TGF)-β-dependent manner^50,51^, the effect of *Cldn11* knockout on TGF-β expression in Sertoli cells and Treg function should be clarified in the future to further understand how CLDN11 prevents autoantibody production.

Permeability barriers against ions and small molecules, but not macromolecules, are reportedly disrupted in MDCK II cells lacking CLDN-based TJ strands^10^. Similar barrier defects are observed in *Cldn5*-deficient mouse endothelial cells with size-selective loosening of the blood–brain barrier^52^. Consistent with those observations, we found that permeability barriers against small molecules (Sulfo-NHS-LC-Biotin: 557 Da) were disrupted in seminiferous epithelia of *Cldn11*^-/-^ mice. Interestingly, however, IgG (approximately 150 kDa), which could not diffuse across normal SCTJs from the interstitium, accumulated at the adluminal compartment of seminiferous tubules in *Cldn11*^-/-^ mice, suggesting that CLDN11 can form permeability barriers against IgG. This is supported by our results that flux of 150 kDa dextran from the interstitium into the adluminal region was increased in *Cldn11*^-/-^ mice. JAM1 is suggested to control macromolecule permeability barriers in the absence of CLDN-based TJ strands in MDCK II cells^10^. Since JAM1 is mislocalized in seminiferous tubules of *Cldn11*^-/-^ mice, CLDN11 may regulate permeability barriers against IgG through JAM1. Taking into consideration that *Cldn11* knockout disrupts physical barriers to IgG, penetration of autoantibodies against antigens in spermatocytes/spermatids into the adluminal compartment of seminiferous tubules may be a cause of defective spermatogenesis in *Cldn11*^-/-^ mice via humoral autoimmune responses. Intriguingly, however, *Rag2* knockout severely impairs the adaptive immune system but did not restore defective spermatogenesis on a *Cldn11* knockout background. Thus, we concluded that CLDN11-mediated spermatogenesis occurs independently of suppressing humoral and cell-mediated autoimmune responses to spermatocytes/spermatids. This is partially supported by our observations that spermatogenesis was impaired without leukocyte infiltration into the seminiferous tubules of *Cldn11*^-/-^ mice. Electron microscopy shows that while some junctional complexes between myoid cells surrounding seminiferous tubules exhibit a continuous interspace of 20 nm, the majority display TJs that effectively prevent the passage of perfused lanthanum into seminiferous epithelia^13^. Thus, myoid cells may act as a semi-permeable barrier to prevent the entry of leukocytes into seminiferous tubules in *Cldn11*^-/-^ mice.

Gow *et al*., also investigated whether CLDN11 plays a key role in immunological barrier formation using *Cldn11*^-/-^ mice^17^, although their data are not shown. In contrast to our results, they have described that sera derived from *Cldn11*^-/-^ mice do not label sections from wild-type testes, and mouse autoantibodies are not detected in the lumina of seminiferous tubules using sheep anti-mouse secondary antibodies to label sections of testes from *Cldn11*^-/-^ mice^17^. These results indicate that autoantibodies against testis-specific antigens are not present in the blood of *Cldn11*^-/-^ mice. Their conclusion could be drawn since autoantibodies were not detected in all sera from *Cldn11*^-/-^ mice, as shown by our immunohistochemistry. Gow *et al*., also found no evidence of CD4-positive cell infiltrates in testes of *Cldn11*^-/-^ mice^17^, which is consistent with our results. Although there is dissimilarity between our and the Gow *et al*., findings associated with autoantibody production in *Cldn11*^-/-^ mice, the spermatogenic phenotypes they observed in *Cldn11*^-/-^ mice (smaller testis size and aberrant spermatogenesis)^17^ are basically similar to those we observed.

The last issue to discuss is how CLDN11 regulate spermatogenesis in mice. We found a significant decrease in the population of differentiating spermatogonia in testes of *Cldn11*^-/-^ mice, which is consistent with previous results^18^, and suggests that CLDN11 is necessary for maintenance of differentiating spermatogonia. A key factor for preserving differentiating spermatogonia is the KIT-mediated signaling pathway. KIT is a tyrosine kinase receptor that is widely used as a marker for differentiating spermatogonia. It is reported that intravenous injection of anti-KIT monoclonal antibody (ACK2), as an antagonistic blocker for KIT function, into adult mice causes depletion of differentiating spermatogonia without affecting undifferentiated spermatogonia^53^. Similar defects in proliferation of differentiating spermatogonia are observed in mice where the codon for Tyr719, a phosphatidylinositol 3-kinase (PI3K) binding site in KIT, is mutated to Phe^54^. The mutation disrupts PI3K binding to KIT and reduces SCF-induced PI3K-dependent activation of AKT^54^. SCF as a ligand for KIT is also a key molecule to maintain differentiating spermatogonia. A recent study using mice with conditional knockout of *Scf* in Sertoli cells demonstrated that SCF derived from Sertoli cells, but not other somatic cells, is crucial for maintenance of the pool of differentiating spermatogonia^29^. Considering that differentiating spermatogonia are located at the basal compartment of seminiferous tubules, SCF is expected to accumulate at the basal compartment to enhance its interaction efficiency with KIT. Consistent with this idea, SCF was localized at Sertoli cell–Sertoli cell junctions near the basement membrane in seminiferous epithelia lacking spermatogenic cells by busulfan treatment. The Sertoli cell junctional localization of SCF was impaired by *Cldn11* knockout. Thus, it is likely that SCF localizes to the plasma membrane of Sertoli cells at the basal compartment, and that SCF localization is regulated by CLDN11. It has been reported that *ZO1*/*ZO2* double knockout MDCK II cells exhibit epithelial polarity defects characterized by intramembrane diffusion of membrane proteins between the apical and basolateral cell surfaces^10^, although the molecular mechanisms through which ZO1/ZO2 control epithelial polarization remain unclear. Since ZO1 localization is severely disorganized in seminiferous tubules of *Cldn11*^-/-^ mice, it is possible that distribution of SCF on the *Cldn11* knockout Sertoli cell surface is affected by epithelial polarity defects. Indeed, apical and basolateral markers were mislocalized in Sertoli cells of busulfan-treated *Cldn11*^-/-^ mice. These findings support the idea that CLDN11 contributes to Sertoli cell polarization, creating a microenvironment for spermatogonial proliferation at the basal compartment of seminiferous tubules through SCF (Fig. 7e). The vast majority of differentiating spermatogonia were lost in *Cldn11*^-/-^ mice, while the surviving differentiating spermatogonia exhibited normal cell proliferative activity, which allowed them to differentiate into spermatocytes/spermatids.

However, increased apoptosis in spermatocytes and complete loss of elongated spermatids were observed in *Cldn11*^-/-^ mice, suggesting that CLDN11 may also regulate differentiation from differentiating spermatogonia to elongated spermatids independently of maintenance of differentiating spermatogonia. To further understand the role of SCTJs in spermatogenesis, we need to investigate them in terms of CLDN11-mediated Sertoli cell polarization as well as Sertoli cell barrier function in the future.

## Methods

### Animals

*Cldn11*^-/-^ mice on a C57BL/6 background were provided from M. Furuse (National Institute for Physiological Sciences, Okazaki, Japan)^34^. To generate *Cldn11*^-/-^/*Rag2*^-/-^ mice, *Cldn11*^-/-^ mice and *Rag2*^-/-^/*Jak3*^-/-^ mice were crossed on a C57BL/6 background (obtained from the Center for Animal Resources and Development, Kumamoto University, Japan; ID576)^41^. *Cldn11*^-/-^ mice and *Cldn11*^-/-^/*Rag2*^-/-^ mice were compared with littermates or age-matched non-littermates from the same colony. Wild-type mice (C57BL/6), WBB6F1-+/+ mice, and WBB6F1-*Sl*/*Sl^d^*mice were purchased from Japan SLC, Inc., (Hamamatsu, Japan). Unless specified otherwise, male mice were analyzed at 10–12 weeks of age. To deplete endogenous spermatogenic cells, mice were intraperitoneally treated with busulfan (44 mg/kg; #B7973; LKT Laboratories). Five weeks after busulfan treatment, testes were collected and fixed. All animals were housed under a 12-h dark– light cycle (light from 7:00 to 19:00) at 22 ± 1°C with *ad libitum* food and water. The Animal Care and Use Committee of Kumamoto University approved the protocols for animal experiments (approval IDs: A2021-016 and A2023-126).

### Cell culture and expression vectors

L cells, MDCK II cells, and human embryonic kidney (HEK) 293 cells were provided by Masatoshi Takeichi (RIKEN Center for Biosystems Dynamics Research, Kobe, Japan), Masayuki Murata (Tokyo Institute of Technology, Yokohama, Japan), and Akira Tsuji (Kanazawa University, Kanazawa, Japan), respectively. Cells were cultured in Dulbecco’s modified Eagle medium (#05919; Nissui) supplemented with 10% fetal bovine serum (FBS) at 37°C under 5% CO_2_. Total RNA was purified from mouse testis using NucleoSpin RNA (#740955.10; Takara). A cDNA library was synthesized from total RNA using ReverTra Ace qPCR RT Kit (#FSQ-101; Toyobo). cDNA encoding mouse CLDN11 (aa 1–207: NCBI accession number NM_008770) and a transmembrane form of mouse SCF (aa 1–245: NCBI accession number NM_001347156) were amplified by polymerase chain reaction (PCR) with KOD-Plus-Ver.2 DNA polymerase (#KOD-211; Toyobo) using a mouse testis cDNA library as a template. For expression of CLDN11 and SCF in mammalian cultured cells, amplified DNA fragments were digested with *Not*I/*Xho*I and subcloned into *Not*I/*Xho*I-digested pCAGGSneodelEcoRI^55^ with or without DNA sequence encoding a C-terminal HA tag. The following primers were used for amplification of cDNA: Forward 5′-AAAGCGGCCGCAGATGGTAGCCACTTGCCTTCAGGTGG-3′, Reverse 5′-AAACTCGAGTTAGACATGGGCACTCTTGGCATGCG-3′ for CLDN11; Forward 5′-AAAGCGGCCGCAGATGAAGAAGACACAAACTTGGATTATC-3′, Reverse 5′-AAACTCGAGCCACCTCTTGAAATTCTCTCTC-3′ for SCF. For bacterial expression, cDNA encoding full-length mouse CLDN11 was amplified by PCR using KOD-Plus-Ver.2 DNA polymerase, digested with *Sal*I/*Not*I and subcloned into *Sal*I/*Not*I-digested pGEX-6p-1 (#28-9546-48; GE Healthcare). The following primers were used for amplification of cDNA: Forward 5′-AAAGTCGACTCATGGTAGCCACTTGCCTTCAGGTGG-3′, Reverse 5′-AAAGCGGCCGCTTAGACATGGGCACTCTTGGCATGCG-3′. To produce GST-fused proteins of the C-terminus of CLDN11 (aa 179–207), expression vectors were constructed by inverse PCR using pGEX-6p-1, in which DNA fragments encoding full-length CLDN11 were inserted into a multicloning site, as a template. The following primers were used for inverse PCR: Forward 5′-TCCGGGGATGCACAGTCATTTGGAG-3′, Reverse 5′-GAGTCGACCCGGGAATTCCGGGGATC-3′. To produce GST-fused proteins of the C-terminus of CLDN11 lacking HV sequence (aa 179–205), expression vectors were constructed by inverse PCR using pGEX-6p-1, in which DNA fragments encoding CLDN11 (aa 179–207) were inserted into a multicloning site, as a template. The following primers were used for inverse PCR: Forward 5′-TAAGCGGCCGCATCGTGACTGACTG-3′, Reverse 5′-GGCACTCTTGGCATGCGTTGGCGAG-3′. Expression vectors for MBP-fused proteins of PDZ1, PDZ2, and PDZ3 domains of mouse ZO1 were kindly provided by Masahiko Itoh (Dokkyo Medical University, Tochigi, Japan)^56^. DNA transfection into mammalian cultured cells was performed using Lipofectamine LTX Reagent with Plus Reagent (#15338-030; Thermo Fisher Scientific), according to the manufacturer’s instructions. Stable cell lines were selected by treatment with 500 μg/ml G-418 (#09380-86; Nacalai Tesque).

### Antibodies

The following primary antibodies were used: rat monoclonal anti-ZO1 (R26.4C; Developmental Studies Hybridoma Bank)^57^; mouse monoclonal anti-ZO1 (T8-754)^58^; rat monoclonal anti-OCLN (MOC37)^59^; rabbit polyclonal anti-OCLN^59^; rabbit polyclonal anti-CLDN3 (#34-1700; Thermo Fisher Scientific); rabbit polyclonal anti-CLDN5 (#34-1600; Thermo Fisher Scientific); rabbit polyclonal anti-CLDN11^32^; rabbit polyclonal anti-CLDN11 (#ab53041; Abcam); rabbit polyclonal anti-JAM1 (#36-1700; Thermo Fisher Scientific); rat monoclonal anti-CDH1 (#M108; Takara); rat monoclonal anti-NECTIN2 (#ab16912; Abcam); rabbit polyclonal anti-GJA1 (#C6219; Sigma–Aldrich); rabbit polyclonal anti-EZR (#sc-20773; Santa Cruz Biotechnology); mouse monoclonal anti-ATP1A1 (#NB300-146; Novus Biologicals); goat polyclonal anti-GFRA1 (#AF560; R&D Systems); rabbit polyclonal anti-PLZF (#HPA001499; Sigma–Aldrich); goat polyclonal anti-LIN28A (#AF3757; R&D Systems); goat polyclonal anti-KIT (#AF1356; R&D Systems); rabbit polyclonal anti-SCP3 (#ab15093; Abcam); rabbit polyclonal anti-VASA (#ab13840; Abcam); rabbit monoclonal anti-WT1 (#ab89901; Abcam); rabbit polyclonal anti-SCF (#ab64677; Abcam; characterized in Supplementary Fig. 13d, e); rabbit polyclonal anti-Ki67 (#ab15580; Abcam); rat monoclonal anti-CD45 (#103101; BioLegend); rabbit polyclonal anti-HA (#561; MBL); and mouse monoclonal anti-α-tubulin (#T6199; Sigma–Aldrich). To obtain serum, blood was collected from mouse heart, incubated for 30 min at room temperature and centrifuged at 2000 × *g* for 20 min. The following secondary antibodies and detection reagents were used: Alexa Fluor 488-conjugated donkey anti-rabbit IgG (#ab150073; Abcam); Alexa Fluor 555-conjugated donkey anti-rabbit IgG (#ab150074; Abcam); Alexa Fluor 488-conjugated donkey anti-mouse IgG (#ab150105; Abcam); Alexa Fluor 594-conjugated donkey anti-mouse IgG (#ab150108; Abcam); Alexa Fluor 488-conjugated donkey anti-rat IgG (#ab150153; Abcam); Alexa Fluor 594-conjugated donkey anti-rat IgG (#ab150156; Abcam); Alexa Fluor 488-conjugated donkey anti-goat IgG (#ab150129; Abcam); Alexa Fluor 594-conjugated donkey anti-goat IgG (#ab150132; Abcam); Alexa Fluor 594-conjugated streptavidin (#S32356; Thermo Fisher Scientific); Alexa Fluor 488-conjugated PNA lectin (#L21409; Thermo Fisher Scientific); horseradish peroxidase (HRP)-conjugated goat anti-rabbit IgG (#ab6721; Abcam); and HRP-conjugated goat anti-mouse IgG (#ab6789; Abcam).

### Histology and immunohistochemistry

Testes were fixed with 4% paraformaldehyde (PFA) in 0.1 M phosphate buffer (PB) at 4°C overnight, dehydrated through an ethanol series, and then embedded in paraffin. Paraffin sections (4 μm thickness) were stained with hematoxylin and eosin according to standard procedures. For immunohistochemistry, sections were subjected to heat-induced antigen retrieval (20 mM Tris-HCl buffer [pH 9.0], 95°C, 15 min). For immunofluorescence staining for IgG, testes were fixed with 4% PFA in 0.1 M PB at 4°C overnight, embedded in OCT compound (#4583; Sakura Finetek), and frozen in liquid nitrogen. Frozen testes were cut into 8-μm thick sections with a cryostat at −20°C. Frozen sections were mounted on microscope slides (#83-1881; Matsunami) and air-dried for 30 min. For immunofluorescence staining using rabbit polyclonal antibodies against TJ-associated transmembrane proteins (OCLN, JAM1, CLDN3, CLDN5, and CLDN11), fresh testes were directly embedded in OCT compound, frozen in liquid nitrogen, and cut into 8-μm thick sections with a cryostat at −20°C. Frozen sections were mounted on coverslips, air-dried for 30 min, fixed in 95% ethanol on ice for 30 min, and treated with 100% acetone for 1 min at room temperature. All sections were washed with phosphate buffered saline (PBS) and blocked with 5% BSA/10% FBS in PBS containing 0.1% Tween 20 for 1 h at room temperature. Primary antibodies and sera (1:300) were diluted in blocking solution and incubated with sections at 4°C overnight. After three PBS washes, sections were incubated with secondary antibodies for 1 h at room temperature. After three PBS washes, samples were mounted in FluorSave Reagent (#345789; Millipore). TUNEL assay was performed using ApopTag Fluorescein In Situ Apoptosis Detection Kit (#S7110; Sigma–Aldrich), according to the manufacturer’s instructions.

For immunofluorescence microscopy of cultured cells, L cells, MDCK II cells, and HEK293 cells were seeded as follows: L cells, 7.5 × 10^5^ cells on coverslips in a 35 mm dish; MDCK II cells, 1.0 × 10^5^ cells on Transwell polycarbonate filters with 12 mm diameter (#3401; Corning); HEK293 cells, 3.0 × 10^5^ cells on coverslips in a 35 mm dish. Two days after cell seeding, L cells were fixed with 1% formaldehyde in PBS containing 0.5 mM CaCl_2_ for 10 min at room temperature. At 3–4 days after cell seeding, MDCK II cells were fixed with 4% formaldehyde in PBS containing 0.5 mM CaCl_2_ for 10 min at room temperature. The day after cell seeding, HEK293 cells were transfected with an expression vector. Two days after transfection, HEK293 cells were fixed with 4% formaldehyde in PBS containing 0.5 mM CaCl_2_ for 10 min at room temperature. After three PBS washes, all samples were permeabilized with 0.2% Triton X-100 in PBS for 10 min at room temperature. Coverslips and filters excised with scalpels were blocked with 1% BSA in PBS for 30 min at room temperature. Primary antibodies were diluted with blocking solution and incubated with samples for 30–60 min at room temperature. After three PBS washes, samples were incubated with secondary antibodies for 30–60 min at room temperature. After three PBS washes, samples were mounted in FluorSave Reagent. All samples were observed using a fluorescence microscope (BX53; Olympus) equipped with a DP74 camera (Olympus) or a confocal laser scanning microscope (FV3000; Olympus). Image acquisition was performed using cellSens Standard software in BX53 or FV31S-SW software in FV3000. Image processing was performed using Fiji/ImageJ (ver. 2.9.0/1.53t; National Institutes of Health).

### Whole-mount immunofluorescence of seminiferous tubules

Testes were collected from mice and transferred into PBS. The tunica albuginea was removed, and the seminiferous tubules teased apart. The interstitial cells were removed by incubating the tubules in 1 mg/ml collagenase (#C0130; Sigma–Aldrich) for 5 min at room temperature. The tubules were then washed with PBS three times and fixed with 1% PFA (for ZO1) or 4% PFA (for KIT) in PBS containing 2 mM CaCl_2_ for 5 h at 4°C. The tubules were washed with PBS three times and permeabilized with 0.25% Triton X-100 in PBS containing 0.05% Tween 20 for 25 min at room temperature. The tubules were again washed with PBS three times and blocked with 5% BSA/10% FBS in PBS containing 0.1% Tween 20 for 1–2 h. Primary antibodies were diluted with blocking solution and incubated with the tubules at 4°C overnight. The tubules were washed with PBS three times and incubated with secondary antibodies for 1–2 h at room temperature. The tubules were washed with PBS three times and mounted in FluorSave Reagent. Samples were observed using FV3000, and Z-stacked images were obtained. Images were processed using Fiji/ImageJ to generate maximum intensity projection views.

### Bead preparation and transplantation

Affi-Gel blue beads (#153-7302; Bio-Rad) were soaked in a solution of BSA (0.5 mg/ml; #A2153; Sigma–Aldrich) or recombinant mouse SCF protein (0.5 mg/ml; #579708; BioLegend) for 1 h at room temperature, as described previously^44^ but with modifications. To distinguish seminiferous tubules adjacent to the transplanted beads, the beads were immersed in 0.83 mg/ml CM-DiI (#C7000; Thermo Fisher Scientific) for 15 min at room temperature. CM-DiI-labeled beads were transplanted into the interstitium of the testes in *Cldn11*^-/-^ mice using syringes with 29G needles (#326666; BD). At 10 days after bead transplantation, testes were collected from mice. The seminiferous tubules were obtained and subjected to whole-mount immunofluorescence staining.

### Tracer assay

PBS containing 2 mM CaCl_2_ and a tracer (EZ-link Sulfo-NHS-LC-Biotin [10 mg/ml; #21335; Thermo Fisher Scientific] as reported previously^40^ or CF488A-conjugated 150 kDa fixable dextran [10 mg/ml; #80131; Biotium]) was prepared. A volume of 15 or 10 μl of solution was injected into the interstitium of testes harvested from *Cldn11*^+/-^ or *Cldn11*^-/-^ mice, respectively, using syringes with 29G needles. After incubation for 30 min on ice, testes were fixed with 4% PFA in 0.1 M PB at 4°C overnight. Testes were dehydrated through an ethanol series and embedded in paraffin to obtain sections.

### Transmission electron microscopy

Testes were fixed by immersion in 2% PFA and 2.5% glutaraldehyde in 0.1 M PB (pH 7.4) for 4 h at 4°C. After washing in 0.1 M PB at 4°C, specimens were postfixed in 1% osmium tetroxide for 2 h at 4°C. They were then washed repeatedly in distilled water, stained with 1% uranyl acetate for 30 min, dehydrated through graded ethanol series and propylene oxide, and embedded in Quetol-812 (#341-H; Nissin EM). Ultrathin sections were cut and mounted onto nickel grids, stained with 1% uranyl acetate for 10 min, followed by Reynolds lead citrate for 5 min, and then examined under a HT7700 electron microscope (Hitachi High-Tech).

### Western blotting

Testes were homogenized on ice in radioimmunoprecipitation assay buffer containing protease inhibitor cocktail (#08714-04; Nacalai Tesque). Homogenates were centrifuged at 15,000 × *g* for 20 min at 4°C. Supernatants were collected, mixed with equal volumes of 2× Laemmli sample buffer supplemented with 200 mM dithiothreitol (DTT), and boiled for 5 min at 100°C. Protein extracts were separated by sodium dodecyl sulfate– polyacrylamide gel electrophoresis (SDS–PAGE) using 12.5% or 15% polyacrylamide gels. Separated proteins were electrotransferred to Immobilon-P polyvinylidene difluoride membranes with 0.45-μm pore size (#IPVH00010; Millipore). Membranes were blocked with 5% skim milk in Tris-buffered saline (TBS) containing 0.1% Tween 20 (TBS-T) for 1 h at room temperature. Primary antibodies were diluted with blocking solution and incubated with membranes at 4°C overnight. Membranes were washed with TBS-T three times and incubated with HRP-conjugated secondary antibodies for 1 h at 37°C. After washing with TBS-T three times, signals were detected by chemiluminescence using Immobilon Western Chemiluminescent HRP Substrate (#WBKLS0100; Millipore), and images were captured using the ChemiDoc Touch Imaging System (Bio-Rad). Image processing was performed using Fiji/ImageJ.

### GST pull-down assay

For *in vitro* binding assays, GST-fused or MBP-fused proteins were expressed in *Escherichia coli* (BL21) (#230280; Agilent) or *Escherichia coli* (DH5α) (#DNA-913F; Toyobo), respectively. Protein expression was induced by addition of 0.3 mM isopropyl β-D-1-thiogalactopyranoside into bacterial culture media, and then bacteria were cultured for 3 h at 37°C. Bacteria were centrifuged at 4,000 × *g* for 15 min. Pellets were resuspended with chilled buffer A (50 mM Tris-HCl [pH8.0], 50 mM NaCl, 1 mM EDTA, and 1 mM DTT), sonicated on ice, and centrifuged at 15,000 × *g* for 25 min at 4°C. Supernatant fluids containing GST-fused proteins or MBP-fused proteins were incubated with Glutathione Sepharose 4B beads (#17-0756-01; Cytiva) or amylose resin (#E8021; New England Biolabs), respectively, for 1 h at 4°C. MBP-fused proteins were eluted from amylose resin by incubation with buffer A containing 10 mM maltose. Eluted MBP-fused proteins were incubated with the Glutathione Sepharose 4B beads coupled with GST-fused proteins at 4°C overnight. Beads were washed three times with buffer A, mixed with equal volumes of 2× Laemmli sample buffer supplemented with 200 mM DTT, and boiled for 5 min at 100°C. Proteins in the supernatant were separated by SDS–PAGE using 12.5% polyacrylamide gels, and gels were subjected to Coomassie Brilliant Blue staining.

### *In silico* data analysis

Single-cell RNA-seq data using spermatogenic cells from adult humans and adult mice were obtained from https://doi.org/10.17632/kxd5f8vpt4.133. Gene expression levels were re-analyzed using Loupe Browser (ver. 6.5.0; 10x Genomics).

### Statistical analysis

In this study, “*n*” refers to the number of mice used for analyses. Data for statistical analyses are shown as mean ± SD or box-and-whisker plots, in which a line in the box shows the median, whiskers represent variation of data, and outliers are plotted as individual points beyond the whiskers. Two-tailed unpaired Student’s *t*-test was performed using Microsoft Excel (ver. 16.78). Dunnett’s test, Mann–Whitney *U* test, Fisher’s exact test, and Tukey–Kramer test were performed using RStudio (ver. 2023.03.1+446; Posit PBC). A *P*-value < 0.05 was considered statistically significant.

## Data availability

Data regarding this paper are available from the corresponding author upon request. Source data are provided with the paper.

## Acknowledgements

We thank Masatoshi Takeichi, Masayuki Murata, Akira Tsuji, and Masahiko Itoh for kindly providing materials; and Kazuya Yoshinaga (Kumamoto University, Kumamoto, Japan), Shosei Yoshida (National Institute for Basic Biology, Okazaki, Japan), Ryusho Kariya (Kobe Gakuin University, Kobe, Japan), and all members of the Wakayama laboratory for discussions and comments. We thank Rachel James, PhD, from Edanz (https://jp.edanz.com/ac) for editing a draft of this manuscript. This work was supported by a Japan Society for the Promotion of Science Grant-in-Aid for Early-Career Scientists (21K15333), Tasaki Memorial Research Grants for 2021 and 2022, the Takeda Science Foundation and Uehara Memorial Foundation (to T.S.).

## Author contributions

T.S. designed the study. T.S., K.S., N.C., K.N., and T.W. performed the experiments. S.O. and M.F. contributed key materials. T.S. and T.W. wrote the manuscript. All authors read and commented on the manuscript.

## Competing interests

The authors declare no competing interests.

## Notes

### Competing Interest Statement

The authors have declared no competing interest.

## References

1. Tsukita, S., Furuse, M. & Itoh, M. Multifunctional strands in tight junctions. Nat. Rev. Mol. Cell Biol. 2, 285–293 (2001).

2. Van Itallie, C. M. & Anderson, J. M. Architecture of tight junctions and principles of molecular composition. Semin. Cell Dev. Biol. 36, 157–165 (2014).

3. Zihni, C., Mills, C., Matter, K. & Balda, M. S. Tight junctions: from simple barriers to multifunctional molecular gates. Nat. Rev. Mol. Cell Biol. 17, 564–580 (2016).

4. Otani, T. & Furuse, M. Tight Junction Structure and Function Revisited. Trends Cell Biol. 30, 805–817 (2020).

5. Farquhar, M. G. & Palade, G. E. Junctional complexes in various epithelia. J. Cell Biol. 17, 375–412 (1963).

6. Staehelin, L. A. Further observations on the fine structure of freeze-cleaved tight junctions. J. Cell Sci. 13, 763–786 (1973).

7. Furuse, M., Fujita, K., Hiiragi, T., Fujimoto, K. & Tsukita, S. Claudin-1 and −2: novel integral membrane proteins localizing at tight junctions with no sequence similarity to occludin. J. Cell Biol. 141, 1539–1550 (1998).

8. Furuse, M., Sasaki, H., Fujimoto, K. & Tsukita, S. A single gene product, claudin-1 or −2, reconstitutes tight junction strands and recruits occludin in fibroblasts. J. Cell Biol. 143, 391–401 (1998).

9. Itoh, M. et al. Direct binding of three tight junction-associated MAGUKs, ZO-1, ZO-2, and ZO-3, with the COOH termini of claudins. J. Cell Biol. 147, 1351–1363 (1999).

10. Otani, T. et al. Claudins and JAM-A coordinately regulate tight junction formation and epithelial polarity. J. Cell Biol. 218, 3372–3396 (2019).

11. Furuse, M. et al. Occludin: a novel integral membrane protein localizing at tight junctions. J. Cell Biol. 123, 1777–1788 (1993).

12. Martìn-Padura, I. et al. Junctional adhesion molecule, a novel member of the immunoglobulin superfamily that distributes at intercellular junctions and modulates monocyte transmigration. J. Cell Biol. 142, 117–127 (1998).

13. Dym, M. & Fawcett, D. W. The blood-testis barrier in the rat and the physiological compartmentation of the seminiferous epithelium. Biol. Reprod. 3, 308–326 (1970).

14. Russell, L. Observations on rat Sertoli ectoplasmic (’junctional’) specializations in their association with germ cells of the rat testis. Tissue Cell 9, 475–498 (1977).

15. O’Rand, M. G. & Romrell, L. J. Appearance of cell surface auto- and isoantigens during spermatogenesis in the rabbit. Dev. Biol. 55, 347–358 (1977).

16. Tung, P. S. & Fritz, I. B. Specific surface antigens on rat pachytene spermatocytes and successive classes of germinal cells. Dev. Biol. 64, 297–315 (1978).

17. Gow, A. et al. CNS myelin and sertoli cell tight junction strands are absent in Osp/claudin-11 null mice. Cell 99, 649–659 (1999).

18. Kanatsu-Shinohara, M., Ogonuki, N., Matoba, S., Ogura, A. & Shinohara, T. Autologous transplantation of spermatogonial stem cells restores fertility in congenitally infertile mice. Proc. Natl. Acad. Sci. U. S. A. 117, 7837–7844 (2020).

19. Chakraborty, P. et al. Androgen-dependent sertoli cell tight junction remodeling is mediated by multiple tight junction components. Mol. Endocrinol. 28, 1055–1072 (2014).

20. Saitou, M. et al. Complex phenotype of mice lacking occludin, a component of tight junction strands. Mol. Biol. Cell 11, 4131–4142 (2000).

21. Shao, M., Ghosh, A., Cooke, V. G., Naik, U. P. & Martin-DeLeon, P. A. JAM-A is present in mammalian spermatozoa where it is essential for normal motility. Dev. Biol. 313, 246–255 (2008).

22. Sakaguchi, T. et al. Putative “stemness” gene jam-B is not required for maintenance of stem cell state in embryonic, neural, or hematopoietic stem cells. Mol. Cell. Biol. 26, 6557–6570 (2006).

23. Russell, L. Movement of spermatocytes from the basal to the adluminal compartment of the rat testis. Am. J. Anat. 148, 313–328 (1977).

24. Smith, B. E. & Braun, R. E. Germ cell migration across Sertoli cell tight junctions. Science 338, 798–802 (2012).

25. Ohta, H., Yomogida, K., Dohmae, K. & Nishimune, Y. Regulation of proliferation and differentiation in spermatogonial stem cells: the role of c-kit and its ligand SCF. Development 127, 2125–2131 (2000).

26. Copeland, N. G. et al. Mast cell growth factor maps near the steel locus on mouse chromosome 10 and is deleted in a number of steel alleles. Cell 63, 175–183 (1990).

27. Zsebo, K. M. et al. Stem cell factor is encoded at the Sl locus of the mouse and is the ligand for the c-kit tyrosine kinase receptor. Cell 63, 213–224 (1990).

28. Huang, E. et al. The hematopoietic growth factor KL is encoded by the Sl locus and is the ligand of the c-kit receptor, the gene product of the W locus. Cell 63, 225–233 (1990).

29. Peng, Y. J. et al. Sertoli cells are the source of stem cell factor for spermatogenesis. Development 150, dev200706 (2023).

30. Flanagan, J. G., Chan, D. C. & Leder, P. Transmembrane form of the kit ligand growth factor is determined by alternative splicing and is missing in the Sld mutant. Cell 64, 1025–1035 (1991).

31. Wehrle-Haller, B. & Imhof, B. A. Stem cell factor presentation to c-Kit. Identification of a basolateral targeting domain. J. Biol. Chem. 276, 12667–12674 (2001).

32. Morita, K., Sasaki, H., Fujimoto, K., Furuse, M. & Tsukita, S. Claudin-11/OSP-based tight junctions of myelin sheaths in brain and Sertoli cells in testis. J. Cell Biol. 145, 579– 588 (1999).

33. Hermann, B. P. et al. The Mammalian Spermatogenesis Single-Cell Transcriptome, from Spermatogonial Stem Cells to Spermatids. Cell Rep. 25, 1650–1667 (2018).

34. Kitajiri, S. et al. Compartmentalization established by claudin-11-based tight junctions in stria vascularis is required for hearing through generation of endocochlear potential. J. Cell Sci. 117, 5087–5096 (2004).

35. Flickinger, C. J. The postnatal development of the Sertoli cells of the mouse. Z. Zellforsch. Mikrosk. Anat. 78, 92–113 (1967).

36. Nagano, T. & Suzuki, F. The postnatal development of the junctional complexes of the mouse Sertoli cells as revealed by freeze-fracture. Anat. Rec. 185, 403–417 (1976).

37. Meng, J., Holdcraft, R. W., Shima, J. E., Griswold, M. D. & Braun, R. E. Androgens regulate the permeability of the blood-testis barrier. Proc. Natl. Acad. Sci. U. S. A. 102, 16696–16700 (2005).

38. Morrow, C. M. et al. Claudin 5 expression in mouse seminiferous epithelium is dependent upon the transcription factor ets variant 5 and contributes to blood-testis barrier function. Biol. Reprod. 81, 871–879 (2009).

39. Nagafuchi, A., Shirayoshi, Y., Okazaki, K., Yasuda, K. & Takeichi, M. Transformation of cell adhesion properties by exogenously introduced E-cadherin cDNA. Nature 329, 341–343 (1987).

40. Chen, Y., Merzdorf, C., Paul, D. L. & Goodenough, D. A. COOH terminus of occludin is required for tight junction barrier function in early Xenopus embryos. J. Cell Biol. 138, 891–899 (1997).

41. Ono, A. et al. Comparative study of human hematopoietic cell engraftment into BALB/c and C57BL/6 strain of rag-2/jak3 double-deficient mice. J. Biomed. Biotechnol. 2011, 539748 (2011).

42. Shinkai, Y. et al. RAG-2-deficient mice lack mature lymphocytes owing to inability to initiate V(D)J rearrangement. Cell 68, 855–867 (1992).

43. Izumi, Y. et al. An atypical PKC directly associates and colocalizes at the epithelial tight junction with ASIP, a mammalian homologue of Caenorhabditis elegans polarity protein PAR-3. J. Cell Biol. 143, 95–106 (1998).

44. Uchida, A. et al. In vivo dynamics of GFRα1-positive spermatogonia stimulated by GDNF signals using a bead transplantation assay. Biochem. Biophys. Res. Commun. 476, 546–552 (2016).

45. Bellvé, A. R. et al. Spermatogenic cells of the prepuberal mouse. Isolation and morphological characterization. J. Cell Biol. 74, 68–85 (1977).

46. Meng, J., Greenlee, A. R., Taub, C. J. & Braun, R. E. Sertoli cell-specific deletion of the androgen receptor compromises testicular immune privilege in mice. Biol. Reprod. 85, 254–260 (2011).

47. Chen, F. et al. ARHGEF15 in Sertoli cells contributes to germ cell development and testicular immune privilege. Biol. Reprod. 107, 1565–1579 (2022).

48. Tung, K. S. et al. Egress of sperm autoantigen from seminiferous tubules maintains systemic tolerance. J. Clin. Invest. 127, 1046–1060 (2017).

49. Barrachina, F. et al. Regulatory T cells play a crucial role in maintaining sperm tolerance and male fertility. Proc. Natl. Acad. Sci. U. S. A. 120, e2306797120 (2023).

50. Fallarino, F. et al. Therapy of experimental type 1 diabetes by isolated Sertoli cell xenografts alone. J. Exp. Med. 206, 2511–2526, (2009).

51. Campese, A. F. et al. Mouse Sertoli cells sustain de novo generation of regulatory T cells by triggering the notch pathway through soluble JAGGED1. Biol. Reprod. 90, 53 (2014).

52. Nitta, T. et al. Size-selective loosening of the blood-brain barrier in claudin-5-deficient mice. J. Cell Biol. 161, 653–660 (2003).

53. Yoshinaga, K. et al. Role of c-kit in mouse spermatogenesis: identification of spermatogonia as a specific site of c-kit expression and function. Development 113, 689– 699 (1991).

54. Blume-Jensen, P. et al. Kit/stem cell factor receptor-induced activation of phosphatidylinositol 3’-kinase is essential for male fertility. Nat. Genet. 24, 157–162 (2000).

55. Niwa, H., Yamamura, K. & Miyazaki, J. Efficient selection for high-expression transfectants with a novel eukaryotic vector. Gene 108, 193–199 (1991).

56. Itoh, M. et al. Junctional adhesion molecule (JAM) binds to PAR-3: a possible mechanism for the recruitment of PAR-3 to tight junctions. J. Cell Biol. 154, 491–497 (2001).

57. Stevenson, B. R., Siliciano, J. D., Mooseker, M. S. & Goodenough, D. A. Identification of ZO-1: a high molecular weight polypeptide associated with the tight junction (zonula occludens) in a variety of epithelia. J. Cell Biol. 103, 755–766 (1986).

58. Itoh, M., Yonemura, S., Nagafuchi, A., Tsukita, S. & Tsukita, S. A 220-kD undercoat-constitutive protein: its specific localization at cadherin-based cell-cell adhesion sites. J. Cell Biol. 115, 1449–1462 (1991).

59. Saitou, M. et al. Mammalian occludin in epithelial cells: its expression and subcellular distribution. Eur. J. Cell Biol. 73, 222–231 (1997).

